# Cryo-tomography reveals rigid-body motion and organization of apicomplexan invasion machinery

**DOI:** 10.1101/2022.04.23.489287

**Authors:** Long Gui, William J. O’Shaughnessy, Kai Cai, Evan Reetz, Michael L. Reese, Daniela Nicastro

## Abstract

The apical complex is a specialized collection of cytoskeletal and secretory machinery in apicomplexan parasites, which include the pathogens that cause malaria and toxoplasmosis. Its structure and mechanism of motion are poorly understood. We used cryo-FIB-milling and cryo-electron tomography to visualize the 3D-structure of the apical complex in its protruded and retracted states. Averages of conoid-fibers revealed their polarity and unusual nine-protofilament arrangement with associated proteins connecting and likely stabilizing the fibers. Neither the structure of the conoid-fibers nor the architecture of the spiral-shaped conoid complex change during protrusion or retraction. Thus, the conoid moves as a rigid body, and is not spring-like and compressible, as previously suggested. Instead, the apical-polar-rings (APR), previously considered rigid, dilate during conoid protrusion. We identified actin-like filaments connecting the conoid and APR during protrusion, suggesting a role during conoid movements. Furthermore, our data capture the parasites in the act of secretion during conoid protrusion.

## Introduction

All intracellular pathogens must accomplish entry into a new host cell. Apicomplexan parasites use an invasion machinery composed of a specialized cytoskeletal complex and secretory organelles. Collectively, this group of organelles is known as the apical complex. It is for this structure that the phylum Apicomplexa is named, which includes the causative agents of malaria, toxoplasmosis, and cryptosporidiosis. Apicomplexan parasites belong to the eukaryotic super-phylum Alveolata, like ciliates and dinoflagellates (Supplemental Figure S1). The apical complex is essential for parasite motility, and for invasion into and egress from host cells. Furthermore, mutations that disrupt the apical complex functions block the Apicomplexan lytic cycle, rendering them non-infectious^1–4^.

The apical complex cytoskeleton itself is a ~250 nm long structure (~25% the length of an *E. coli*) composed of a series of rings organized around a central spiral of specialized tubulin fibers called the conoid, and inside which secretory organelles are organized and prepared for secretion (Figure 1A). Whereas the conoid fibers are composed of the same tubulin dimers that make up the parasites’ subpellicular microtubules, they do not form closed tubes^5^. Instead, the conoid fibers form an open “C”-shaped structure^5^. The conoid complex is highly dynamic and protrudes and retracts^6^ as the parasites secrete adhesins and other motility/invasion factors^7,8^. Remarkably, the apical complex appears to have evolved from a eukaryotic cilium, as it contains both tubulin and cilium-associated proteins^9–12^. In addition, the core of the apical complex, including the conoid and its specialized tubulin structures, is conserved not only throughout Apicomplexa^13,14^ and in closely related free-living organisms^15^ but also in more distantly related Alveolata, such as dinoflagellates^16,17^ (Supplemental Figure S1). Thus the apical complex appears to be an ancient structure, of which the molecular composition, high-resolution structure, and mechanistic understanding of its functions are still largely a mystery.

**Figure 1.**
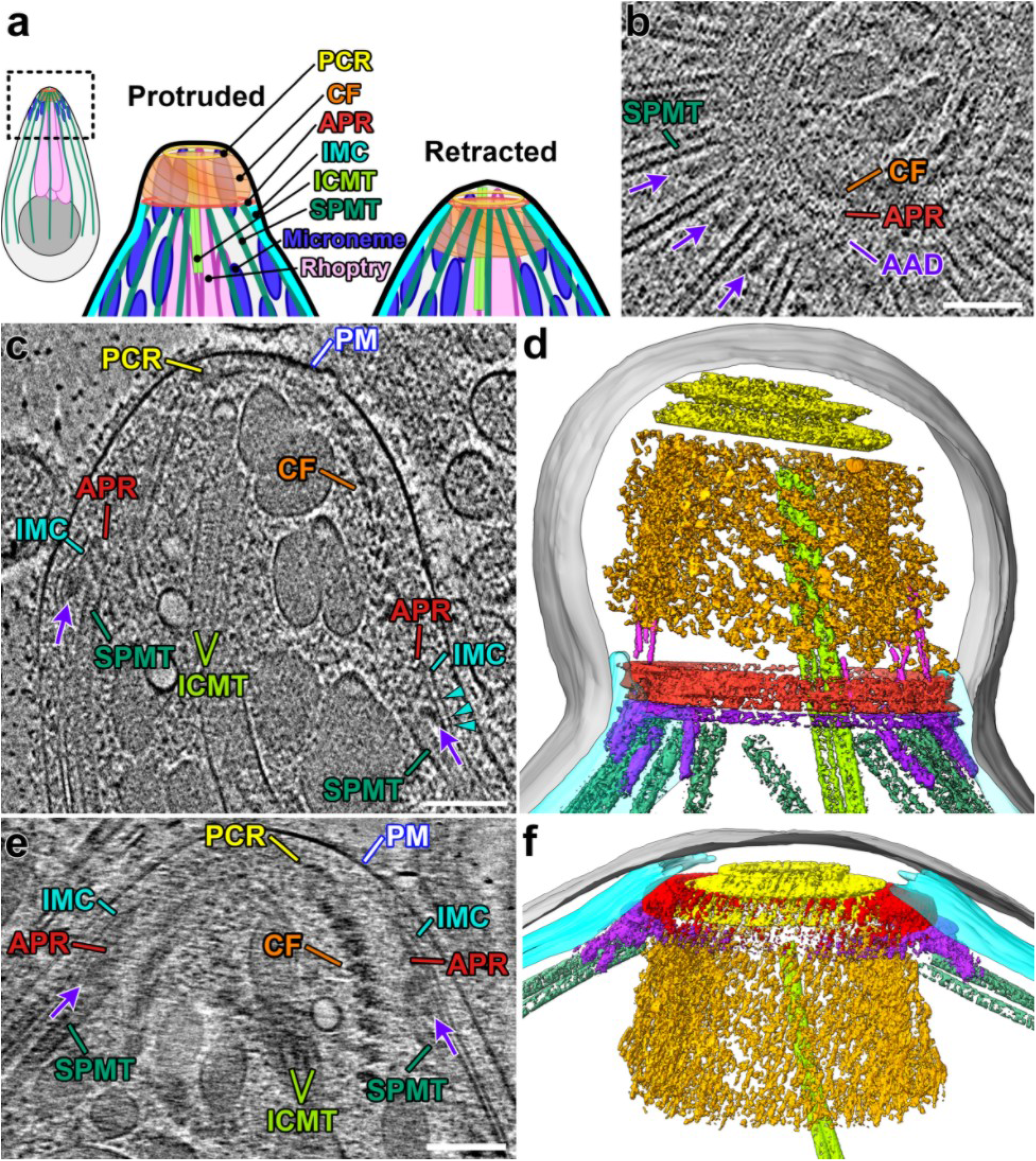
Three-dimensional *in situ* architecture of the apicomplexan invasion machinery revealed by cryo-FIB milling and cryo-ET. **(A)** Cartoon overview of the components of the coccidian apical complex comparing the protruded and retracted states. This coloring scheme will be used throughout the manuscript. **(B)** Tomographic slice through a partially retracted conoid that was cryo-FIB milled in cross-sectional orientation, clearly shows the SPMTs ending near the AAD ring and with AAD projections (purple arrows) interspersed between neighboring SPMTs. **(C)** Tomographic slice of the reconstructed apical end of *N. caninum* with protruded conoid. Note the density connecting between the two membranes of the inner membrane complex (IMC, cyan arrowheads). Other labels and coloring see below. **(D)**3D segmentation and visualization of the apical complex from a protruded conoid (different tomogram from (C)). Labels and colors used throughout the manuscript unless otherwise noted: AAD (purple), amorphous APR-associated density ring and projections; actin-like filaments (magenta in D); APR (red), apical polar rings; CF (orange), conoid fiber; ICMT (light green), intraconoidal microtubules; IMC (cyan), inner membrane complex; PCR (yellow), pre-conoidal rings; PM (gray), plasma membrane; SPMT (dark green), subpellicular microtubules. **(E)** Tomographic slice of the reconstructed apical end of a milled *N. caninum* with a retracted conoid, annotated as in (C). **(F)**3D segmentation and visualization of a retracted conoid (different tomogram from (E)) and colored as in **(D)**. In longitudinal views, apical tip is oriented towards the top of the images throughout the manuscript, unless otherwise noted. Scale bars: 100 nm (in B, C, E).

Common ultrastructural characteristics of the Alveolata super-phylum include the cytoskeleton-supported vesicular structures that lie just basal to the plasma membrane and are known as alveoli^18^. In apicomplexans these structures are called the inner membrane complex (IMC), which runs along the length of the parasite^19–22^. Apicomplexan parasites have two distinct sets of specialized organelles, called the micronemes and rhoptries, from which invasion factors and effector proteins are secreted (Figure 1A)^23–25^. Whereas the rhoptry secretion requires close contact with the host cell plasma membrane, micronemes are thought to secrete continuously while the parasites are extracellular. Although these secretory organelles are not broadly conserved among alveolates, recent work has demonstrated that secretion from the rhoptries is mediated by a complex that is conserved both in its structure and protein components in ciliates such as *Tetrahymena* and *Paramecium^26,27^*.

Cellular cryo-electron tomography (cryo-ET) is a powerful imaging technique that can reveal structural details with molecular resolution, but the suitable thickness of biological samples is limited to a few hundred nanometers^28^. Most intact eukaryotic cells are thicker than this. Therefore, previous cryo-ET studies of whole eukaryotic cells have imaged either naturally thin cell regions, such as lamellipodia^29^ and cilia^30^, or cells that were compressed due to embedding in a thin layer of ice (compression force by water surface tension)^31,32^. Thicker specimens can be made thin enough for cryo-ET either by frozen-hydrated sectioning using a knife – which is prone to cutting artifacts^33^ – or by cryo-focused ion beam milling^34^.

We have applied cryo-ET to the coccidian apicomplexan parasites *Toxoplasma gondii*, arguably the most widespread and successful parasite in the world, and its close relative *Neospora caninum*. Our study used mainly cryo-focused ion beam (cryo-FIB) milling of unperturbed cells embedded in thick ice to compare the coccidian apical complex in its protruded and retracted states *in situ*, without the compression and deformation artifacts that have plagued studies of unmilled intact apicomplexan samples^31,32,35^. Our tomographic reconstructions provide an unrivaled view of the apical complex structure. Subtomogram averaging of the conoid fibers allowed us to clearly demonstrate that, contrary to a popular hypothesis^5,35^, the conoid is not spring-like, and does not deform during cycles of protrusion and retraction. We also observed filaments that extend between the conoid and the apical polar rings (APR) through which the conoid moves, suggesting that polymerization of actin, or an actin-like protein, may play a role during conoid motion. Our data also provide an unprecedented view of the interactions between the parasite’s secretory organelles and the apical complex cytoskeleton. Finally, we were able to capture the act of microneme fusion with the intact plasma membrane.

Together this work provides structural and mechanistic details for the apicomplexan invasion machinery, the apical complex. Because this complex is essential for apicomplexan parasite infection, understanding the molecular basis of its function could reveal important new targets for therapeutic intervention for some of the world’s most devastating diseases.

## Results

### Cryo-FIB-milling followed by cryo-electron tomography preserves the 3D structure of the parasite cytoskeleton

The coccidian cytoskeleton appears remarkably well-preserved after detergent-extraction and negative-staining, which has led to many TEM studies of apicomplexan ultrastructure using this preparation method^1,4,5,36^. We first tested whether cryo-ET of detergent-extracted and then plunge-frozen *Toxoplasma* cells would allow us to gain an accurate 3D view, with improved resolution of the cytoskeletal structures, as compared to conventional TEM of negative-stained (dried) samples. Although cryo-ET of detergent-treated samples reveal cytoskeletal assemblies such as the apical conoid and subpellicular microtubules at high contrast (Supplemental Figure S2), the tomograms also show artifacts from the detergent extraction as well as structural distortions *(e.g*., flattening; compare Supplemental Figure S2E-F to S2G-H) that complicated gaining reliable structural information from these samples.

Therefore, to obtain the least perturbed and highest quality samples, we plunge-froze intact and live parasites in a relatively thick layer of ice (>1 μm thick; to avoid cell flattening by water surface tension). We then used cryo-FIB milling to generate 150-200 nm thick lamella of the vitrified, but otherwise native, parasites (Supplemental Figure S3). To generate these samples, we used the NC1 strain of *Neospora caninum*, which is a BSL1 organism closely related to the human pathogen *Toxoplasma gondii* (Supplemental Figure S1). The conoid of extracellular parasites protrudes and retracts continuously – with and without host cells present, but just before plunge-freezing, the parasites were either prepared in an intracellular-like buffer^37^ or incubated with 10 μM calcium ionophore for 10 min. so that the majority of parasites have conoids preferably in the retracted or protruded states^6^, respectively, without blocking conoid motility or cellular functions like secretion. The resulting cryo-tomograms reveal well-preserved structural details of the native *N. caninum* apical complex, including membranes, cytoskeletal assemblies, and organelles (Figure 1, Movie S1).

### Structural organization of the apical cell region

We compared the architecture of the *N. caninum* apical complex in two states: conoid protruded and retracted. In the protruded state (Figures 1C,D; Movie S1), the pre-conoidal rings (PCRs) are the cytoskeletal structure closest to the apical plasma membrane (Figure 1C,D), followed by the associated conoid that consists of 14-15 conoid fibers arranged in a spiral (Supplemental Figure 2G,H). As compared to the protruded conoid state, the conoid fibers in the retracted state are tucked just basal to the APRs, and the PCRs are located slightly apical of the APR structures (Figure 1E,F). Basal to the protruded conoid, the inner membrane complex (IMC) is clearly visible as a flat double membrane structure with associated densities just below the parasite plasma membrane (Figure 1C,E). Notably, our *in situ* cryo-ET reveals several (non-periodic) densities that bridge between the two membranes of the extended sheet-like IMC (cyan arrowheads in Figures 1C; 2C), perhaps serving as “spacers” that constrain the distance between the membranes. The apical rim of the IMC sheet is attached to the apical polar rings (APRs; Figure 1C-F). Just beneath the IMC, about 22 subpellicular microtubules extend basally from the APRs (Figure 1B-E). Although the subpellicular microtubules and associated microtubule inner proteins (MIPs) are well-resolved in our cryo-tomograms, we have not focused on them here, because they have previously been analyzed in detail^35,38^.

**Figure 2.**
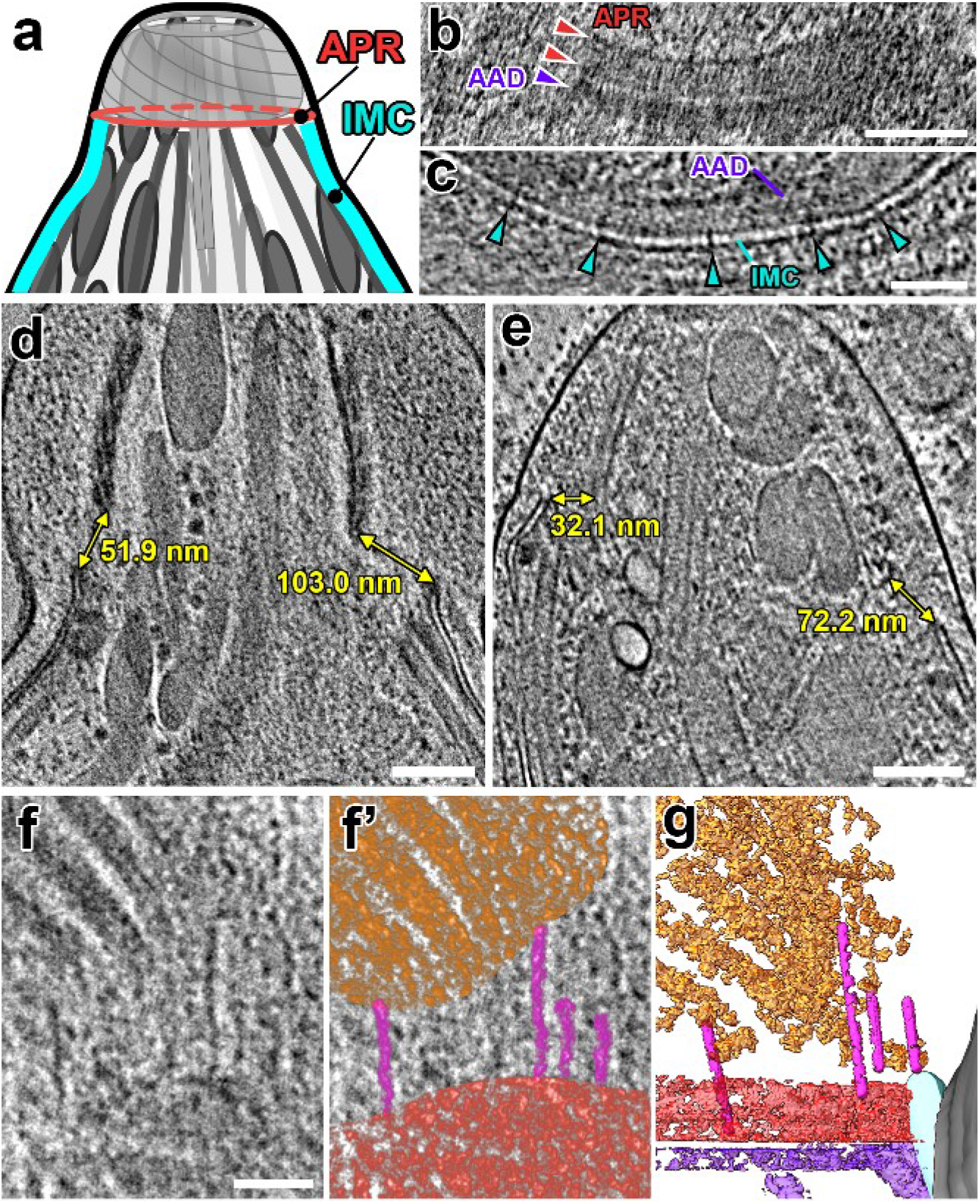
The protruded conoid is tilted and off-center relative to the APRs. **(A)** Cartoon of the apical complex in the protruded state, highlighting the APR and IMC structures. **(B)** A tomographic slice (longitudinal orientation) shows two distinct APR rings (red arrowheads) and the AAD ring (purple arrowhead) which is located between the APR and IMC. **(C)** A crosssectional tomographic slice shows the IMC near the apical edge with “spacer” densities (cyan arrowheads) between the two membranes. **(D, E)** Tomographic slices through the protruded conoid complex from 2 parasites. Measurements mark the minimum distances between the IMC and the conoid fibers on each side of the conoid. **(F-G)** Tomographic slice (F: original; F’: pseudo-colored) and 3D-segmented isosurface rendering (G) of the same region at the base of a protruded conoid show filamentous actin-like densities (magenta) connecting between the conoid (orange) and the APRs (red). Scale bars: 100 nm (in B, D, E); 50 nm (in C, F).

Super-resolution light microscopy has suggested components of the APR segregate into independent rings^39^, but these rings have been difficult to resolve with negative staining EM. In our tomograms, we can clearly resolve several ring-shaped structures near the apical rim of the IMC: two distinct APRs with similar diameters, *i.e*. the A1 ring (apical) and the thicker A2 ring (basal) that may consist of two closely stacked sub-rings A2a and A2b (Figure 2A,B, Supplemental Figure S2D, S4A-C). We also observe a ring of amorphous density (named here “amorphous APR-associated density” or AAD) that is located between the IMC and the tips of the subpellicular microtubules (purple arrows/structure in Figure 1B-F; Supplemental Figures S2A,D, S4A-C). The AAD appears to be connected to the APR rings and has basal projections (21 nm wide and 64 nm long) that are sandwiched between adjacent subpellicular microtubules (Figure 1B). The AAD projections have not been reported from conventional EM studies, but have recently been observed in another cryo-ET study (called “interspersed pillars” in Sun, *et al*.^35^). We observe the APRs, AAD ring, and AAD projections in tomograms of both the protruded and retracted conoid states. Interestingly, the AAD ring and projections appear preserved during detergent-extraction of the parasite (Supplemental Figures S2A,D, S4A-B). This localization and biochemical behavior suggest that the AAD ring and projections may contain components of the IMC apical cap, which have recently been localized by super-resolution light microscopy as interleaved between microtubules^4,40^. However, in these studies, known apical cap proteins such as ISP1 and AC9 extend ~1 μm from the APR, far beyond the ~64 nm occupied by the AAD projections. Therefore, the AAD ring and projections appear to be composed of proteins that have yet to be identified or precisely localized.

Surprisingly, and in contrast to the idea that the APR serves to strongly anchor the conoid to the parasite membrane, the protruded conoid never appears fully square to the APRs in our tomograms (compare protruded in Figures 1C,D; 2D,E, to retracted in Figure 1E,F; Supplemental Figure S4D-F). Instead, the conoid appears able to tilt and be off-center relative to the annular APR as it protrudes. To quantify this observation, we compared the distances between the apical rim of the IMC and the basal edge of the conoid from opposing sides in central slices through our tomograms (Figure 2D,E; Supplemental Figure S4D-F). We found that the difference between the opposing distances varied more in tomograms of the protruded state (Δd = 52±12 nm; n=3), than the retracted state (Δd = 10±4 nm; n=3). These data are consistent with the idea that the conoid is “leashed” to the APRs and/or the apical rim of the IMC with the AAD by flexible and perhaps dynamic structures. Indeed, we frequently observed filaments between the APR region and the conoid fibers (Figures 1D, 2F-G; Supplemental Figure S4G-L).

Previous studies showed that apicomplexan “gliding motility” is driven by treadmilling of actin fibers between the IMC cytoskeleton and the parasite plasma membrane^41,42^. Recently, F-actin nanobodies were used to demonstrate that actin fibers nucleate at the conoid and are passed down the parasite as it moves^3^. We clearly observe filaments consistent in diameter with actin fibers (~8 nm diameter) extending from the conoid to connect with the APR region, near where the hand-off to the IMC-associated myosin network^41,43^ would be expected to occur. These filaments vary in length from 38 – 164 nm in our tomograms (n=14 fibers from 3 tomograms), suggesting they are dynamic in nature (Figures 1D, 2F-G; Supplemental Figure S4G-L).

In contrast to intact parasites, in our detergent-extracted samples, the conoid base appears to sit directly on the APRs (Supplemental Figure S2A, S4A,B), as is typically seen in similarly treated samples by negative staining EM^4,5,36^. This collapse of the gap and filamentous structures incorrectly suggests that the APR and its connections to the conoid are much more rigid than we observe in our cryo-FIB-milled, but otherwise unperturbed, samples, which again highlights the value of our *in situ* analysis of the structures in their native states.

### Conoid structure is unchanged during cycles of protrusion and retraction

As described below, the conoid fibers are unusual tubulin polymers in which the protofilaments form an open C-shaped cross-section (Movie S1). These fibers were initially described as wound in a “spring-like” structure^5^, leading to a model that the conoid motions were driven by a spring-like mechanism, involving cycles of deformation in the conoid structure followed by release. This hypothesis is made more attractive because of the conoid fibers unusual C-shape, which would be expected to be more deformable/compressible than closed microtubules. Prior to the development of cryo-FIB milling, electron microscopy and tomography analysis of the conoid required detergent extraction of the parasite membrane and/or cell flattening, which often result in structural artifacts during sample preparation and during image processing, where orientation bias prevents proper missing-wedge correction (Supplemental Figure S2A-F,I). Our data of the unperturbed, native apical complex allows us to unambiguously address changes of the conoid ultrastructure during different functional sates.

We therefore sought to assess whether the conoid movements behaved according to the spring-like model involving deformation of the conoid and conoid fibers. We reasoned that any gross ultrastructural change in the conformation of the conoid polymer would result in changes in the overall dimensions and architectures of the conoid. We therefore compared the dimensions of protruded and retracted conoids in tomograms from cryo-FIB milled *N. caninum* (Figure 3; Supplemental Figure S5). Neither the apical conoid diameters (244 ± 19 nm protruded vs. 236 ± 8 nm retracted), the basal diameters (350 ± 25 nm vs. 326 ± 7 nm), nor the conoid heights (277 ± 5 nm vs. 262 ± 13 nm) were significantly different between the two states (Figure 3E; Supplemental Figure S5A). Similarly, the angle of the conoid fibers relative to the PCRs, and the intervals between adjacent conoid fibers were indistinguishable in the two states (Figure 3F-I). Furthermore, both the mean conoid fiber length and the distribution of lengths in the population were not significantly different between the protruded and retracted states (399 ± 37 nm vs. 405 ± 45 nm; Supplemental Figure S6A).

**Figure 3.**
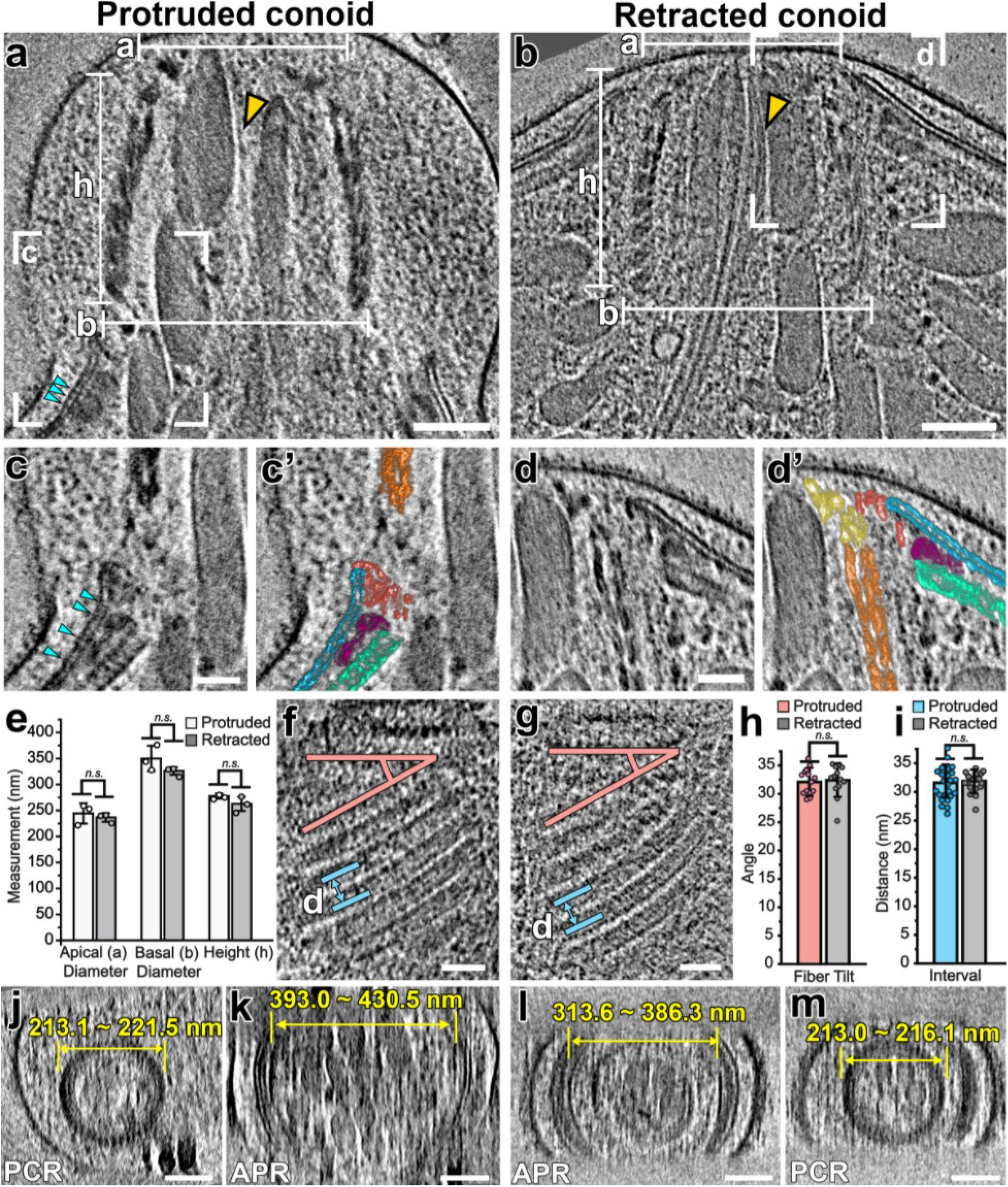
The overall structure of the conoid remains unchanged during the cycles of protrusion and retraction. **(A-D’)** Tomographic slices through the center of the apical complex show the conoid in the protruded (A) and retracted (B) states. White lines indicate the measurements of the apical diameter (a), the basal diameter (b) and the height (h) of the conoid structure. White boxes in (A, B) indicate the areas magnified in (C, D) where the conoid and the APR-associated complex interact; (C’ and D’) are the pseudo-colored versions of (C and D), colored as detailed in Figure 1. Note the elongated, sheet-like density (gold arrowheads in A and B) tracking between micronemes and ICMT inside the conoid. IMC “spacer” densities are indicated with cyan arrowheads in (A,C). **(E)** Measurements of the apical diameter, the basal diameter and the height of protruded (white bars) and retracted (gray bars) conoids show no significant changes during the cycles of protrusion and retraction. **(F-I)** Tomographic slices (F, G) through the edge of reconstructed conoids (showing the conoid fibers in longitudinal section) in the protruded (F) and retracted (G) states. Lines indicate the measurement of the relative angle between the CFs and the PCR plane (red lines), and the measurement of the distance between neighboring CFs (blue lines); the results of latter measurements for protruded and retracted conoids are shown in (H) and (I), respectively. **(J-M)** Tomographic slices show representative PCRs (J and M) and APRs (K and L) in cross-sectional views of the apical complex in protruded (J,K) and retracted (L,M) states. Diameters of APRs and PCRs are indicated as ranges from all available tomograms (n=3 for APRs in both states; n=3 PCRs protruded; n=2 PCRs retracted). Scale bars: 100 nm (in A,B,J-M); 50 nm (in C,D,F,G). Statistical significance was calculated by Student’s t-test. n.s., not significant (p > 0.05).

Whereas the apical tip of the conoid and the PCRs are level with the APR when the conoid is retracted, after extension of the conoid, the base of the conoid is located in the region of the APR (compare Figure 3A,C,C’ with 3B,D,D’). In switching between the protruded and retracted states, the apical IMC regions, the AAD, and the APRs all change their position relative to the conoid and its associated PCRs, and to the plasma membrane (Figure 3A-D’). The positions of the APRs and AAD appear closely coupled to the apical rim of the IMC in our tomograms, which appears to flex apically in the protruded state. We therefore asked whether the APRs exhibited structural changes correlated with conoid protrusion and retraction. We measured the diameters of the APRs in each of our tomograms and determined that there was a striking 10~30% increase in APR diameter in the protruded versus retracted states (Figure 3K,L; Supplemental Figure S5B). In contrast to the APRs, the structure of the PCRs and conoid showed no observable differences between the two states (Figure 3J,M; Supplemental Figure S5C).

### Molecular structure and polarity of the conoid fibers

We next assessed the conoid fiber structure at a higher resolution by generating separate subtomogram averages of the conoid fibers from the protruded (Figure 4A; ~3.0 nm resolution at 0.5 FSC, see Supplemental Figure S6B) and retracted states (Figure 4B; ~3.2 nm resolution at 0.5 FSC, see Supplemental Figure S6B). Visually, the subtomogram averages of the conoid fibers from either state appear largely indistinguishable, and reveal the 9 tubulin protofilaments^5^, as well as distinct protein densities decorating the fibers (Figure 4A,B). To better assess any variation between the two states, we fitted tubulin dimers into the two averaged cryo-ET maps using Chimera^44^ (Figure 4C,D) with correlation-coefficients of ~0.9. We then overlaid the best-fit models of a single repeat of the conoid fiber from each state (Figure 4E). Consistent with the lack of difference between the shape of the entire conoid structure between the two states (Figure 3), each of the 9 protofilament subunits appear positioned indistinguishably between the two states within the resolution of the data and error of the fit (Figure 4E). Note that the three protofilament subunits with the highest RMSD in their overlay (PF5-7; Supplemental Figure S6C) are fitted in a lower resolution region of the averages.

**Figure 4.**
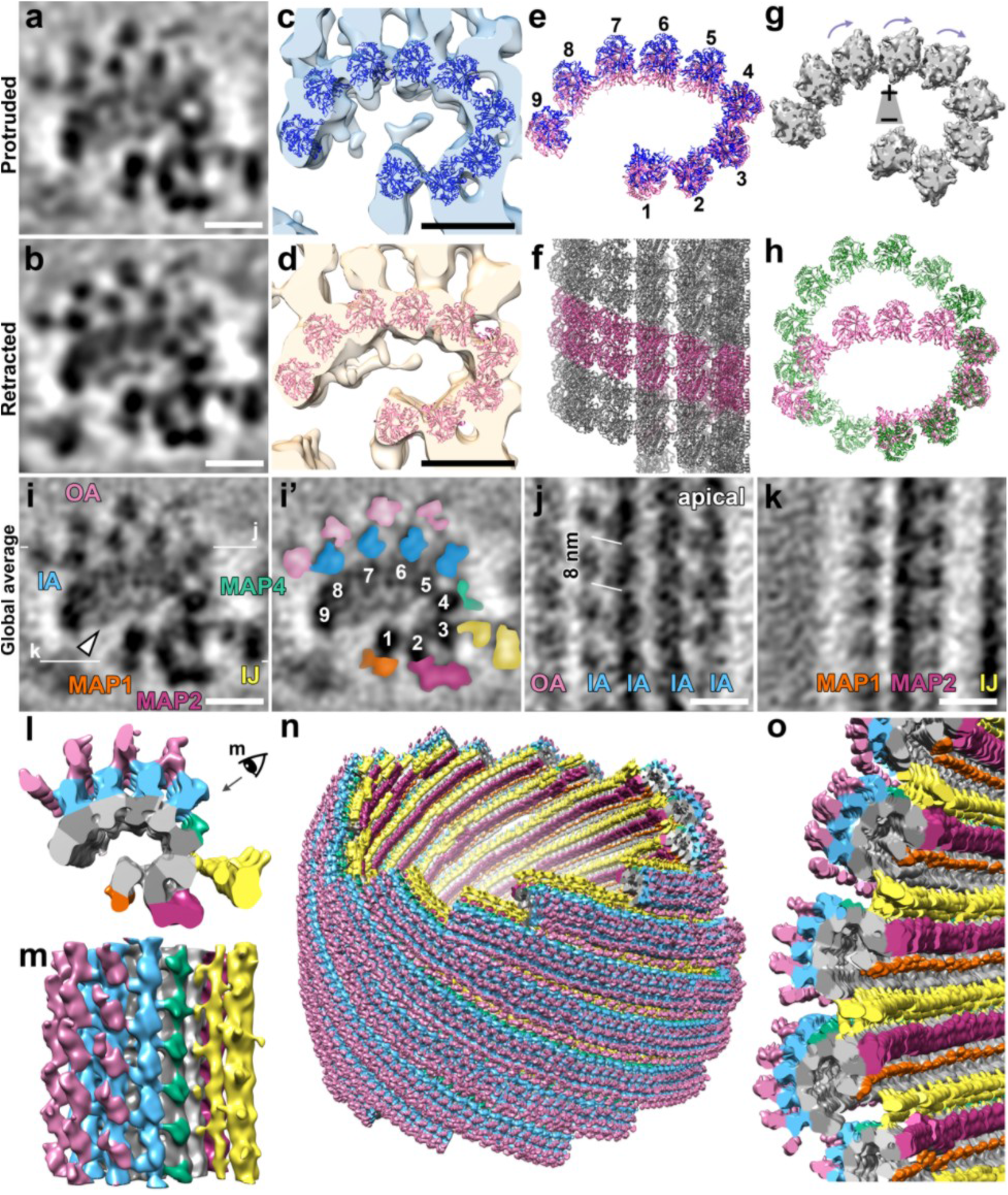
Subtomogram averages of the conoid fibers show a C-shape architecture with nine tubulin protofilaments and associated proteins. **(A, B)** Cross-sectional tomographic slices of the averaged 8-nm repeats of the CF fibers in the protruded (A) and retracted (B) states. **(C-E)** The high-resolution structure of tubulin was fitted into the subtomogram averages of the protruded (C, blue) and the retracted (D, pink) states. Comparison (E) of the two pseudo-atomic protofilament models in the protruded (blue) and retracted (pink) states shows no significant difference. **(F)** Longitudinal views of the pseudo-atomic protofilament model in the retracted state. The pitch was estimated based on the rise of the periodic CF-associated MAPs between neighboring protofilaments. **(G)** Isosurface rendering of the nice-fold averaged protofilaments display a ‘clockwise skew’ when viewed from the conoid base, suggesting the minus ends of the conoid fibers are orientated to the base. **(H)** The arrangements of the modeled protofilaments in the CFs (pink) compared with the high-resolution cryo-EM structure of the typical 13-protofilament subpellicular microtubules (green; EMDB: EMD-23870). The most substantial difference is the angle change between PF4 and PF5, causing a tight kink in the protofilament arrangement. **(I-K)** Tomographic slices of the global CF subtomogram averages that combine all data from both the protruded (A) and retracted (B) states viewed in cross-sectional (I: original; I’: pseudo-colored) and longitudinal (J and K) orientations. The white lines in (I) indicate the locations of the slices in the respective panels. Labels and coloring see below. **(L-O)** Isosurface renderings show the 3D structures of the averaged CF repeats in cross-sectional (L) and longitudinal (M) views, as well as the CFs from a complete conoid (N) by assembling the averaged 8-nm repeats back into the full tomogram. This and the zoom-in (O) show that the open face of the C-shaped CFs faces the interior of the conoid. Labels and coloring: 1-9, protofilaments; IA (blue) and OA (pink), “inner-layer arm” and “outer-layer arm” densities; IJ (yellow), inner junction MAP; MAP1 (orange), MAP2 (magenta), MAP4 (green), microtubule associated proteins. Scale bars: 10 nm (in A, B, I-K).

Taken together, our data demonstrate that the conoid appears to move as a rigid body during protrusion and retraction, rather than as a conformationally deformable structure, and there is no obvious rearrangement of protofilaments within the conoid fibers during this process. From these data, we cannot, however, rule out minor conformational changes between the two states. Our *in situ* data also corroborate that the conoid fibers form an unusual open C-shaped arrangement of 9 protofilaments^5^ that requires contacts distinct from a typical 13-protofilament MT (Figure 4H).

Having demonstrated that the conoid fibers do not undergo major conformational changes between the protruded and retracted states, we combined particles from both states to calculate a global average of the conoid fibers (Figure 4I-K). The resulting average of the two states shows markedly improved signal-to-noise ratio and a resolution of ~2.8 nm at the 0.5 FSC criterion (or ~2.2 nm at the 0.143 FSC criterion; Supplemental Figure S6B) and reveals density coating both the external (MT-associated proteins; MAPs) and internal (MT inner proteins; MIPs) edges of the conoid fiber (Figure 4I-O; Movie S1). The MIP densities bridge multiple protofilaments and coat almost the entire inner surface of the conoid fiber (Figure 4I,I’). These contacts may help explain the apparent rigidity of the conoid fiber structure. The external surface of the conoid fibers also appears almost entirely decorated with MAPs (Figure 4I-O; Supplemental Figure S2I-K). Associated with PF5-8 are two layers of MAPs, named here as “inner-layer arm” and “outer-layer arm” densities (IA and OA; Figure 4I-J; Supplemental Figure S2I-J). Additional MAPs include MAP1, 2, and 4 which are associated with PF1, 2, and 4 respectively. We also observe inner-junction MAP densities (IJ) that connect PF3 of one conoid fiber (n) with PF9 of the neighboring fiber (n+1; Figure 4I,I’,K-O; Supplemental Figures S2I,I’,K, S6F-H). These inter-fiber contacts likely stabilize the spiral conoid structure and reinforce its rigidity. Most MAPs exhibit a clear 8 nm periodicity along the length of the conoid fiber (Figure 4J,K,M; Supplemental Figure S2J-K, S6E,E’).

Regular microtubules are helical assemblies that have a characteristic handedness, helical pitch, and polarity. For example, a typical 13-protofilament microtubule forms a straight polymer with a rise of 3 monomers (12 nm) per left-handed helical turn (a “13-3 helix”), or 0.92 nm rise per protofilament-to-protofilament contact^45^. Using the regularly spaced MAPs as a guide, we were able to assign the pitch of the conoid fiber in longitudinal views (Figure 4F,J,K; Supplemental Figures S2J, S6E,E’). As the C-shaped conoid fibers are formed from an open, rather than closed-helical microtubule, we calculated only the protofilament-to-protofilament pitch. Our model indicates that the conoid fiber forms through a left-handed assembly – like a typical microtubule, but has an estimated pitch of ~1.5 nm protofilament-to-protofilament, which is ~1.6-fold larger than a typical microtubule (Figure 4F; Supplemental Figure S6E,E’). To determine the structural polarity of the conoid fiber, we performed a nine-fold average of the protofilaments (using the tangential/normal to the C-shape curvature of the fiber). The isosurface rendering of the 9-fold averaged protofilaments displayed a ‘clockwise skew’ when viewed from the conoid base, suggesting the minus ends of the conoid fibers are located at the base of the conoid complex (Figure 4G).

### Subtomogram averages of the pre-conoidal rings reveal 3 layers of periodic densities that are connected

Instead of the previously reported two PCRs^5,9^, we resolve three PCRs (Figure 5; Supplemental Figures S2B; S7): P1 is the most apical and smallest ring (Figure 5B-D,G,H; Supplemental Figure 7E,I) with 151 ± 6 nm diameter – which was likely overlooked in previous detergent-treated and negatively stained samples; P2 (middle; Figure 5B-H; Supplemental Figure S7F,J) consists of two subrings with the following diameters: P2a – 178 ± 2 nm, P2b – 217 ± 8 nm; and P3 (basal; Figure 5C,D,G,H; Supplemental Figure S7G,K) has a diameter of 258 ± 8nm (Figure 5B-H). All diameters were measured on the outside edges of the rings in 4 different cells (n=4). Subtomogram averaging of the PCR from four reconstructed *N. caninum* cells produced a 4.9 nm resolution average (0.5 FSC criterion, Supplemental Figure S6B), and revealed both the periodicity and connections between the PCR layers. In our tomograms, each of the layers appear to have the same periodicity, with 45-47 subunits per ring (Figure 5G,H). P2 and P3 are connected with regularly spaced linkers that are ~25 nm long, with apparent bridging density between the neighboring linker subunits (Figure 5B-D,G; Supplemental Figures S2B, S7C).

**Figure 5.**
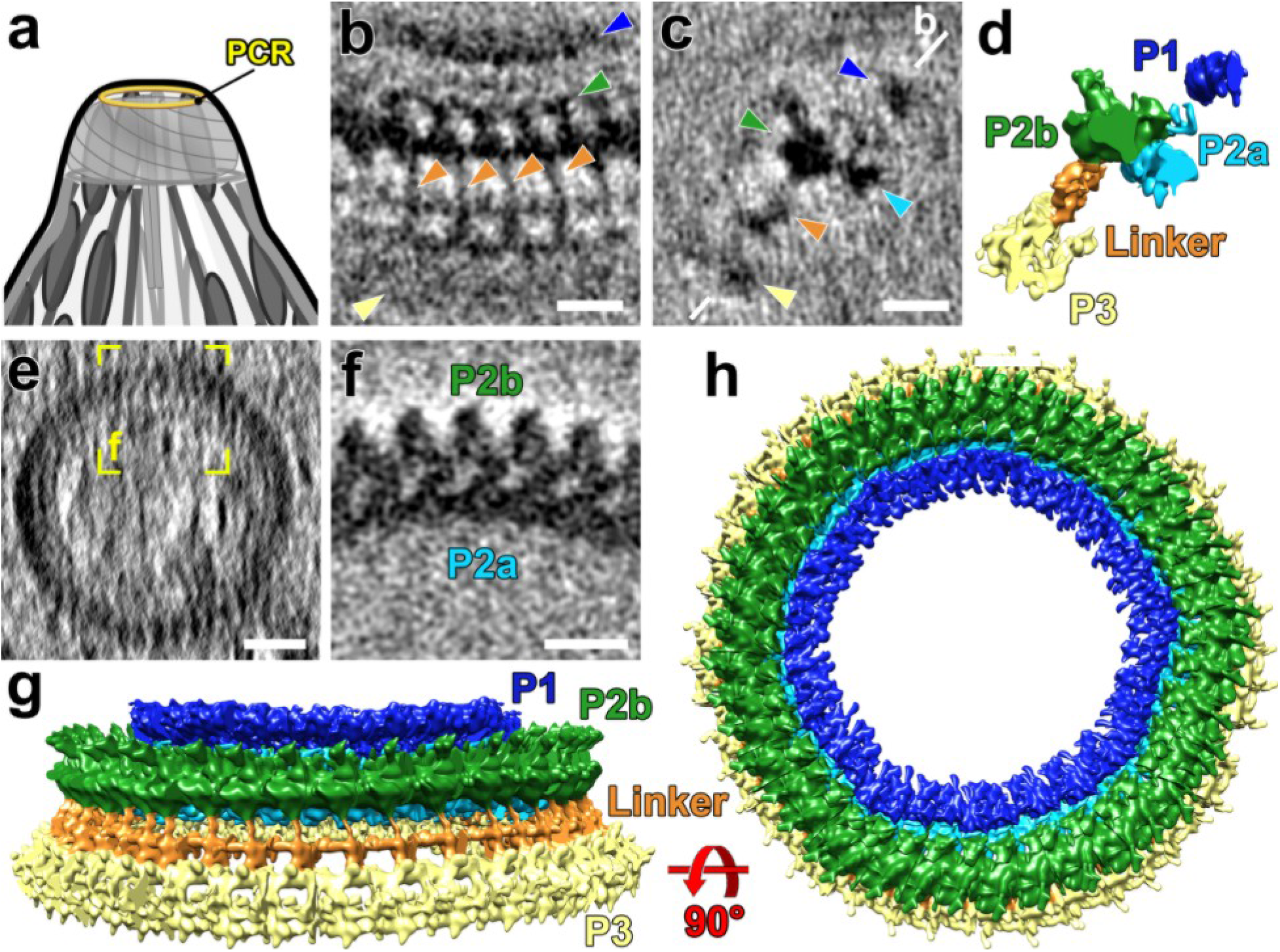
Subtomogram averages of the PCRs show three rings and linker. **(A)** Cartoon of the apical complex highlighting the location of the PCRs. **(B-D)** Tomographic slices (B and C), and isosurface rendering of the averaged PCR repeats (D) viewed in the tangential (B) and longitudinal (C and D) orientations, showing different components of the PCR including the apical P1 (highlighted by blue arrowheads), P2a (cyan), P2b (green), P3 (yellow) and the linker (orange) between P2b and P3. The white line in (C) indicates the location of the slice in (B). **(E-F)** Cross-sectional slices from a raw tomogram (E) and the averaged PCR repeats (F) show that the round PCR P2 ring is composed of an outer and an inner ring, P2a and P2b, respectively. Yellow square in (E) indicates the orientation and location of the subtomogram average displayed in (F). **(G, H)** Isosurface renderings show the complete PCRs by assembling the averaged repeats back to the full tomogram. Scale bars, 20 nm (in B, C, and F); 50 nm (in E).

### In situ cryo-ET of native cells reveals functional segregation of secretory organelles within the coccidian conoid

Conventional EM of chemically-fixed and resin-embedded apicomplexan parasites has provided a basic overview of the organization of the apical secretory organelles within the coccidian conoid^5,46^, which we used to guide our analysis of the *in situ* structure of the apical complex – including the secretory organelles – in our tomograms (Figures 6 and 7; Supplemental Figure S8; Movie S1). We clearly observe the central pair of intraconoidal microtubules (ICMT), extending from the conoid tip basally for 500-800 nm (Figure 6A-B; Supplemental Figure S6I), which is longer than the 350 nm previously reported^5^. The ICMT have been proposed to be a major organizing center for secretion in coccidia^47,48^, and appear to have a set of associated proteins that are distinct from both the subpellicular microtubules and conoid fibers^5^, though only one such protein has been identified to date^49^.

**Figure 6.**
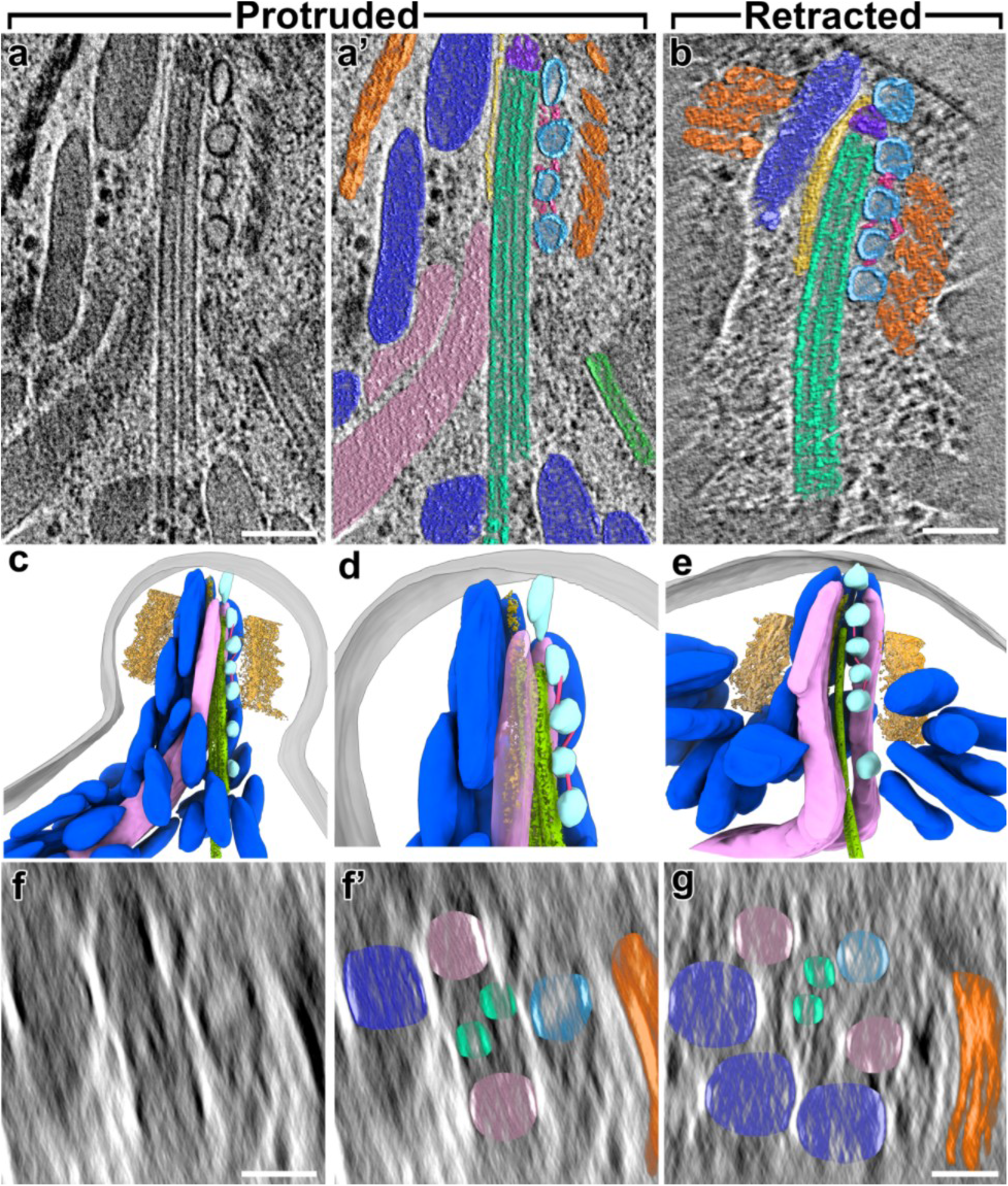
The apical secretory machinery is organized around the intra-conoidal microtubules (ICMT) in both protruded and retracted states. **(A-B)** Tomographic slices show the apical tip of a *N. caninum* cell with the secretory organelles organized around the ICMT (green) inside the protruded conoid complex in longitudinal views (A, A’ – protruded original and pseudo-colored; B – retracted). The long ICMTs connect the apex of the conoid to the cytosol, and closely co-localized with the secretory vesicles (light blue) and two rhoptries (rose). Note that the membrane-associated, most-apical vesicle is not visible in the tomographic slice shown in (A), but is visible in panels (C,D) and Figure 7B. Other coloring: CFs (orange), micronemes (dark blue), sheet-like density (gold), inter-vesicular connections (pink), “crowning” density that caps the apical minus-end of the ICMT (purple). **(C-E)**3D segmentation and visualization of tomograms show in (A and B, respectively), i.e. with protruded (C, D) and retracted (E) conoid, showing the overall organization of secretory organelles within the conoid complex, including CFs (trimmed from the front to show the content inside), micronemes, rhoptries, ICMT, vesicles, inter-vesicular connections, sheet-like structure along micronemes, and plasma membrane (gray). (D) shows a zoom-in from (C), with conoid fibers hidden for clarity. **(F-G)** Cross-sectional slices through the protruded (F, F’ – original and pseudo-colored) and retracted (G) conoids from the tomograms shown in (A and B). Scale bars: 100 nm (in A,B); 50 nm (in F,G).

**Figure 7.**
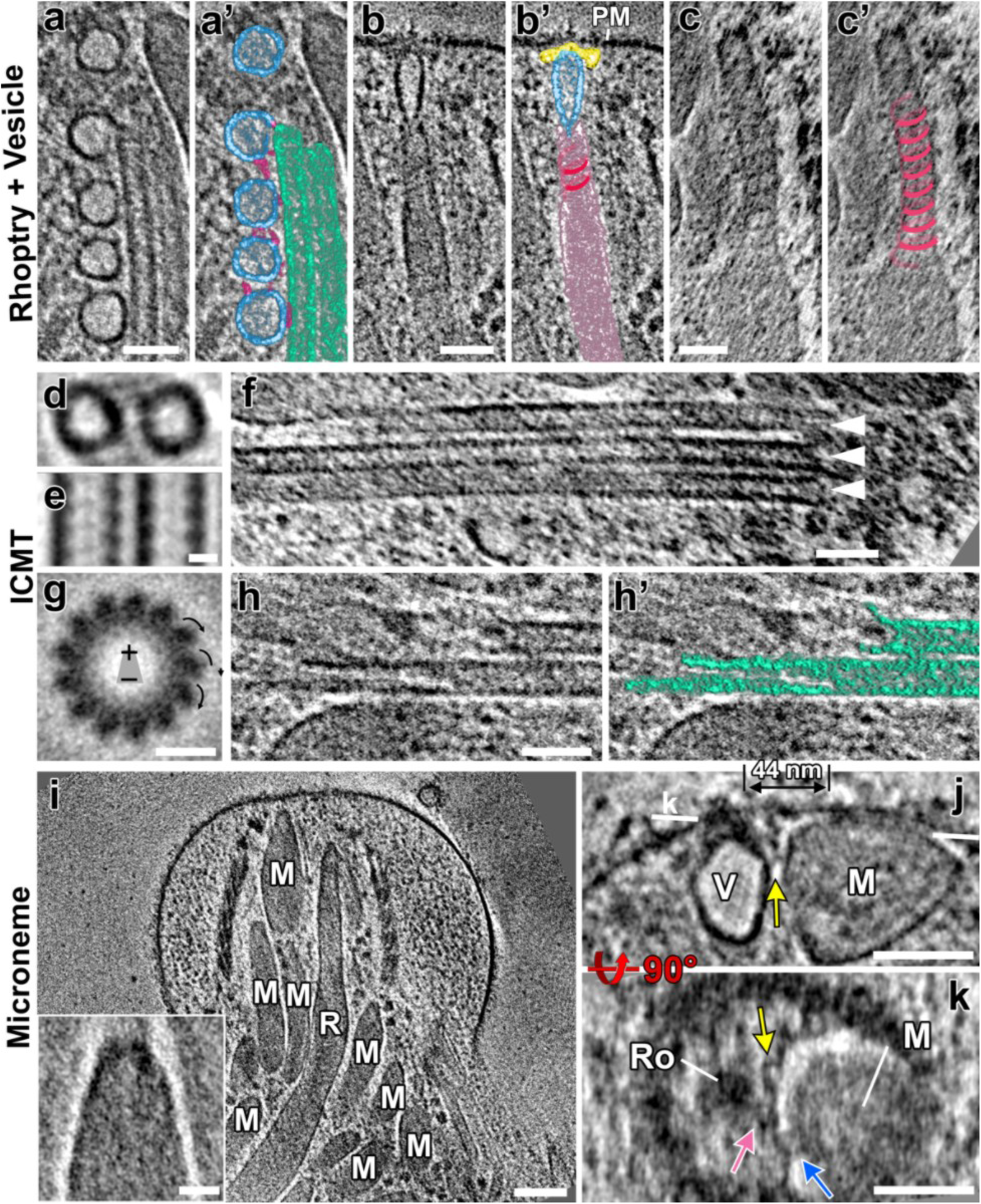
The intra-conoidal microtubules and functional segregation of secretory organelles within the conoid complex revealed by cryo-FIB milling and cryo-ET. **(A-C’)** Tomographic slices (A,B,C: original; A’,B’,C’: pseudo-colored) show: (A, A’) 5 regularly spaced vesicles (light blue), of which 4 are tracking along one of the microtubules of the ICMT (light green) and are connected by inter-vesicular linkers (pink); (B, B’) a rhoptry (rose) interacting with the plasma membrane (PM) via the most-apical vesicle (light blue) and the “rosette” docking complex (yellow); and (C, C’) a spiraling scaffold (dark rose) associated with the rhoptry membrane in the rhoptry-neck region. **(D, E)** Slices of the subtomogram averaged 8-nm repeats of the ICMTs viewed in cross-sectional (D) and longitudinal (E) orientations. **(F)** Occasionally, more than two microtubules were observed in the ICMT complex. Shown here is an example of three microtubules (white arrowheads). **(G)** Thirteen-fold rotationally averaged ICMT from 53 sub-tomograms of detergent-extracted *Toxoplasma* cells. The arrows indicate the clockwise skew of the protofilaments when the ICMT are viewed from apical to basal, indicating that the minus ends of the ICMTs are oriented apically in the parasite. **(H, H’)** The basal ends of the ICMTs showed flared ends and different lengths of protofilaments, which is usually associated with dynamic plus-ends of MTs. **(I)** A tomographic slice of a protruded conoid shows the organization of micronemes. Insert: the subtomogram average of 25 microneme apical tips shows a flattened, electron dense cap. **(J, K)** Tomographic slices provide a side (J) and a top cross-sectional view (K) of two secretory organelles, a microneme (M) and a rhoptry-associated vesicle (V), that are docked side-by-side to the plasma membrane (the vesicle through the rosette (Ro)), but with distinct docking sites (44 nm apart). Note that both organelles are tethered (pink and blue arrows) to the same plasma membrane-anchored ridge (yellow arrows). The white line in (J) indicates the location of the slice in (K). Scale bars: 100 nm (in I); 50 nm (in A-C, F, H, J and K); 20 nm (in I insert), 10 nm (in D, E and G)

In line with what has been observed with EM of resin-embedded samples^5^, we observe that distinct secretory organelles (the rhoptries and micronemes) segregate to opposite sides of the ICMT in tomograms from both protruded and retracted conoids (Figure 6). We also observed 4-6 regularly spaced vesicles tracking along one of the microtubules of the ICMT (Figures 6A-E; 7A,A’), consistent with previous reports^26,46^. We observed a single vesicle just apical of the ICMT that appears docked to the density of the recently described “rosette” (Figure 7B-B’), and which has been associated with facilitating rhoptry secretion^26^. Consistent with this function, we observe that the most-apical, membrane-docked vesicle connects to the apical tips of 1-2 rhoptries (Figure 7B,B’; Supplemental Figure S8A). We also identified clear densities connecting the vesicles to both the ICMT and to each other within a chain (Figures 6A-E; 7A). Notably, we did not observe this bridging density at the apical, membrane-docked vesicle (Figure 7A; n=6 tomograms with the vesicle chain and apical vesicle present; note that in other tomograms the apical vesicle was milled away by cryo-FIB). Whereas the apical vesicle likely originated from the ICMT-associated vesicle chain, in docking with the rhoptry and plasma membrane, it appears to have been released from the vesicle-chain and its connections. We also observed that some of the vesicles appear elongated along the axis of the ICMT, suggesting that they are under tension and have been captured in active trafficking along the ICMT (Figure 6A).

A parasite cell has ~10 rhoptries and tens of micronemes, though only a subset of either organelle is docked at the parasite conoid at any given time^46^. In all tomograms that contained the full diameter of the parasite conoid (6 of 9 tomograms), we observed 2 rhoptries tracking along the ICMT, but on opposite sides of the ICMT (Figure 6C-G; colored rose). However, we do not observe clear linkers connecting the rhoptries to the ICMT. Within the apical region of the parasite, a spiral-shaped density corkscrews along the rhoptry membranes (Figure 7B-C’), reminiscent of the localization pattern of Ferlin-2 seen by immuno-EM^50^. The filaments spiraling up the rhoptries within the conoid were previously estimated to have a diameter of ~33 nm and a pitch of ~21.5 nm from tomograms of unmilled samples in which the parasites had been flattened due to surface tension^51^. We found that such a predicted helix is consistent with our measurements of ~36 nm in diameter and ~17 nm in pitch from cryo-FIB-milled parasites (Figure 7C,C’).

Because of their short length and potential variation in their associated proteins along their length, subtomogram averaging of the ICMT did not yield sufficient resolution to clearly resolve MAPs or MIPs (Figure 7D,E). Instead, we examined the chirality of the ICMT cross-section^52^ using an ICMT subtomogram average from a detergent-extracted tomogram (Figure 7G). This analysis demonstrates that both of the paired ICMT, unlike the conoid fibers, have their minus-ends oriented apically. Moreover, the basal, plus-end of the ICMT pair exhibits the splayed morphology typical of a dynamic microtubule (Figure 7H). This observation suggests that the ICMT, unlike most other apicomplexan MT structures, exhibits dynamic instability at its plus-end (Figure 7H), and may explain why the previously reported ICMT lengths vary largely between samples and are consistently shorter in extracted samples^5^ versus those that we measure from intact, native parasites. We also observe a density that caps the apical, minus-end of the ICMT (Figure 6A-B), which would be consistent with a microtubule-organizing center from which the ICMT are polymerizing. In coccidia, however, γ-tubulin appears to be restricted to the parasite centrioles and cytoplasm, and has not been localized to the apical complex^53^. As has been previously reported^46^, some ICMT contained a third microtubule (Figure 7F), suggesting that rigorous control of the structure is unnecessary for its function.

Notably, the ICMT and its associated vesicles and rhoptries segregate to one side of the conoid that is distinct from the region occupied by the micronemes (Figure 6C-G), thus the ICMT are not directly involved in organizing micronemes for secretion. Nevertheless, the micronemes do not appear disorganized (Figure 6C-E; 7I). Instead, they are arrayed in clusters and show a distinct polarized orientation. The micronemes are relatively uniform in length (220±30 nm) and width (58±11 nm; Supplemental Figure S8B-H), and the basal ends are rounded, whereas the apical ends appear narrow with a flattened, electron-dense cap (Figure 7I; inset). This density suggests an undescribed scaffolding complex that we propose organizes the micronemes and assists in their trafficking. Furthermore, in one tomogram of a retracted conoid, we observed two micronemes that appear to be docked through their apical tips with the plasma membrane (Supplemental Figure S8I). In addition, we consistently identified a long sheet-like density between the micronemes and the ICMT in our tomograms (Figure 6A-B; Movie S1). The sheet varies in thickness between about 4-8 nm, appears fibrous, is about 15-30 nm wide, and can often be traced to extend from the top of the conoid to below its base. These data suggest that the sheet-like structure may assist in the organization and trafficking of the secretory organelles.

Whereas micronemes secrete continuously to promote the motility of extracellular parasites, they are also intimately involved in triggering host cell invasion. To initiate invasion of a host cell, all apicomplexan parasites rely on the secretion of a receptor/co-receptor pair comprising a microneme protein (AMA1) and a rhoptry proteins (RON2)^54,55^. Microneme and rhoptry components of the invasion apparatus have been proposed to mix as they were secreted through a common path at the conoid^23^. That we observe micronemes segregated from the docked rhoptries calls this model into question. Furthermore, in one tomogram of a protruded conoid, we identified a microneme that appears to have been captured in the process of plasma membrane docking/secretion (Figure 7J,K; Supplemental Figure S8J-M). Consistent with an ongoing secretion event, the microneme in question is about half the length of all other micronemes measured (Supplemental Figure S8H, red data points). We observe a density consistent with one large or multiple contact sites in a circular region of ~85 nm in diameter (Figure 7K; Supplemental Figure S8J,L). We also observe an apparent bolus of density on the surface of the plasma membrane associated with the contact site (Figure 7J; Supplemental Figure S8J,M), consistent with secretion of microneme contents. Notably, the docking site of this secreting microneme is 44 nm from the rosette and its docked rhoptry/apical vesicle, indicating that the two organelles secrete at separate sites using distinct secretory machineries (Figure 7J). However, both the rhoptry-associated vesicle and the docked microneme are tethered to the same ridge (Figure 7J-K; Supplemental Figure S8J; yellow arrows) and membrane-anchor (Supplemental Figure S8M and inset; yellow arrowheads), which would likely facilitate spatial and temporal coordination of secretion.

## Discussion

The conoid structure of the apical complex has captured the imagination of microscopists since its earliest descriptions by conventional TEM^56,57^. Here, we have applied cryo-ET to cryo-FIB-milled parasites to compare the native *in situ* structures of the coccidian apical complex cytoskeleton and associated secretory organelles in the protruded and retracted states of the conoid. Conoid protrusion is associated with striking morphological changes at the parasite apical tip that are visible by light microscopy^6,58^. These morphological changes, coupled with the conoid’s unusual spiral shape, has led to a model that the conoid is spring-like and deforms during its movements (protrusion/retraction). Previous cryo-EM of detergent-extracted parasites^59^ and a recently published cryo-ET analysis^35^, document differences in protruded versus retracted conoids. However, subtomogram averaging in the Sun et al. study focused on detergent-extracted samples, and the samples used in both previous analyses were compressed, which can lead to structural artifacts.

In contrast, our cryo-FIB-milled samples preserve the circularity of the coccidian apical complex structure (Supplemental Figure S2G-H; Movie S1), which allows more reliable interrogation of its native structure. Our data show that both the ultrastructure and molecular organization of the protruded and retracted states of the conoid are indistinguishable, indicating the conoid fibers do not deform like a spring during conoid movements, and therefore do not provide energy to assist in either conoid motion or in secretion.

Whereas we did not observe changes in the structure of the retracted versus protruded conoid, we found that the APR appears to dilate during protrusion which appears coupled with flexion of the IMC at the apical tip during protrusion. We also did not observe close contacts between the conoid and the APR, rather the protruded conoid appears often somewhat tilted relative to the APR-plane, consistent with flexible and/or dynamic tethers connecting the conoid to the APR and/or the apical rim of the IMC. RNG2 appears to be one such protein, as its N- and C-termini appear to span between the conoid and APR during protrusion^60^. Whereas we would not expect to identify density of such an intrinsically disordered protein, we did observe densities that span between the base of the protruded conoid and the APR that were consistent with actin/actin-like filaments. Therefore, it is possible that actin polymerization at this site is at least partially responsible for generating the force of conoid motion. Notably, an early description of conoid protrusion found that depolymerization of actin with cytochalasin D attenuated, but did not completely abrogate, conoid protrusion^6^. Recently, Formin-1, which is responsible for polymerization of actin at the conoid, has been demonstrated essential for conoid protrusion^61^. Nevertheless, further high-resolution structural studies using genetic perturbation will be required to place individual proteins in the apical complex and tease apart the physical basis of the conoid dynamics and function.

Because our cryo-FIB-milled samples preserved the parasite membranes, we obtained an unparalleled view of the interactions between the cytoskeleton of the apical complex and the parasite’s specialized secretory organelles. We observed close contacts between the ICMT and both the parasite rhoptries and the apical vesicle chain. This suggests that the ICMT are responsible for organizing these organelles, but not the micronemes, with which we observed no direct contacts. Intriguingly, whereas the ICMT are not broadly conserved in Apicomplexa outside of Eucoccidiorida, a pair of ICMT have been reported in *Chromera velia*^15^, a free-living alveolate that is among the extant organisms most closely related to Apicomplexa (Supplemental Figure S1). Thus, the ICMT were likely present in the ancestral species. That the apical vesicles and rhoptries track along the ICMT suggests that a microtubule motor may be facilitating their organization, though parasite kinesins and dyneins have not been localized to the ICMT. Because the majority of trafficking in Apicomplexa appears to occur on actin filaments^3,62–64^ the parasite microtubule motors have not yet been systematically characterized. Notably, only a single ICMT-localized protein has been identified to date^49^, which lacks clear homology to known structures.

A major open question in the field is the molecular basis of the force that drives the secretion of the invasion-associated organelles docked at the apical complex. The rhoptries are >1 μm in length, and likely require a more intricate machinery than simple membrane docking to drive secretion. Intriguingly, depolymerization of actin using cytochalasin D blocks parasite motility and invasion but not host cell attachment and rhoptry secretion^65^, suggesting actin is not responsible for driving secretion. Recent work identified the apicomplexan Nd proteins that comprise the “apical rosettes”, structures best characterized in the ciliate group of Alveolata^27^. Similar to their role in ciliates, the apicomplexan Nd proteins are essential for rhoptry secretion^26^, and the rosette structure was recently described using cryo-ET of unmilled parasites^51^. The *Toxoplasma* protein Ferlin-2 is also required for rhoptry secretion^50^, and we and others^51^ identified a density spiraling around the rhoptries within the conoid that is reminiscent of published Ferlin-2 immuno-EM staining pattern^50^. These spiraling filaments may act somewhat like dynamin to squeeze the rhoptries during secretion. Among the *Toxoplasma* Nd-associated proteins were putative GTPase-related proteins^26^, suggesting a dynamin-like activity may be present at this site as well.

With the ability to rapidly freeze and thus capture snapshots of highly dynamic cellular processes, we were able to observe a docked microneme that appears to be in the process of secretion. We found that the site of microneme secretion is distinct from that of a docked rhoptry-associated vesicle, suggesting that the components of the invasion machinery secreted by these two organelles must find each other in the apical plasma membrane after secretion. Our data indicate that microneme secretion can occur during conoid protrusion, events that have been indirectly correlated by other studies^7^. Note however, that these data do not rule out additional secretion during the retracted state. Finally, we not only observed a plasma membrane-associated ridge and anchor between the docked microneme and rhoptry-associated vesicle, but also an elongated (fibrous) sheet that tracks between the micronemes and the ICMT. These structures may represent cytoskeletal elements responsible for organization, trafficking and coordinated membrane docking of the apical secretory organelles. Microneme turnover requires trafficking along actin, though biogenesis and organization of micronemes appear actin-independ^63^ent^63^. Further studies will be required to identify the molecular components of the anchor, ridge and sheet, and their functions in organellar trafficking and secretion.

In summary, cryo-ET of cryo-FIB-milled parasites has enabled us to examine the apical complex *in situ*, overcoming the compression artifacts that occur when preparing intact cells in a thin layer of ice. These significant advances allowed us to interrogate the native structure of the conoid complex and its movements during retraction and protrusion. We were able to unambiguously demonstrate that the conoid moves as a rigid body during protrusion, with filamentous, actin-like projections that connect it to the APR. The next frontier in understanding the mechanics of the apical complex will be in capturing parasites during host cell invasion. It is possible that components of the apical complex will undergo some conformational change when the parasite is in intimate contact with a host cell membrane. We also anticipate that further studies combining proteomic data with cryo-ET of cryo-FIB-milled parasites will enable the placement of individual proteins into the apical complex structure, which will shed new light onto the molecular and structural basis of its movements and function.

## Materials and Methods

### Phylogenetic analysis

The phylogenetic tree in Supplemental Figure S1 was estimated in RAxMLv8.2^66^ using the LG substitution model with gamma rate heterogeneity and empirical frequencies with 1000 bootstrap using an alignment of the protein sequences for HSP90 concatenated with RPS11 (accessions: AFC36923.1, BESB_021480, XP_029219085.1, LOC34617734, XP_022591029.1, ETH2_0701200, ETH2_0910900, NCLIV_040880, XP_003884203.1, TGME49_288380, XP_002366350.1, SN3_03000005, SN3_01300510, PBANKA_0805700, XP_034421046.1, PF3D7_0708400, XP_001351246.1, PVP01_0108700, PVP01_0822500, GNI_014030, XP_011128490.1, KVP17_001483, KAH0483594.1, FG379_001268, KAH7649672.1, Chro.30427, OLQ16118.1, cgd3_3770, CPATCC_001922, Vbra_12473, CEM00719.1, Cvel_2184, Cvel_482, AAA30132.1, XP_764864.1, XP_952473.1, XP_952423.1, XP_001611554.1, XP_001609980.1, XP_002775585.1, XP_002766754.1, XP_001447795.1, XP_001445466.1, XP_001009780.1, XP_001030186.1, AAR27544.1, XP_009040431.1, XP_009033899.1).

### Cell culture and cryo-preparation

Human foreskin fibroblasts (HFF) were grown in Dulbecco’s modified Eagle’s medium supplemented with 10% fetal bovine serum and 2 mM glutamine. *Toxoplasma gondii* (RH strain) and *Neospora caninum* (NC1 strain) tachyzoites were maintained in confluent monolayers of HFF. Detergent-extracted *Toxoplasma* cells and the associated subtomogram averages are displayed in Figure 7G and Supplemental Figures S2A-F,I-K, S4A-B, and S6I. Tomographic reconstructions of cryo-FIB-milled *N. caninum* cells, and the corresponding subtomogram averages and data analyses are presented in Figures 1-6, 7A-F,H-K and Supplemental Figures S2G,H, S3, S4C-L, S5, S6A-H, and S7–S8. For preparation of parasites for cryo-ET, highly infected HFF monolayers were mechanically disrupted by passage through a 27 gauge needle to release the parasites. For “retracted conoid” samples, parasites were kept in “Endo Buffer” (44.7 mM K_2_SO_4_, 10 mM MgSO_4_, 106 mM sucrose, 5 mM glucose, 20 mM Tris-H2SO_4_, 3.5 mg/mL BSA, pH to 8.2 with H_2_SO_4_), which preserves the parasites in an intracellular-like state^37^. For “protruded conoid” samples, parasites were kept in HEPES pH 7.4 buffered saline after release from cells. All parasites were passed through a 5 μm filter to remove cell debris washed in appropriate buffer and collected by centrifugation for 10 min at 300 g. Parasites were resuspended in the respective buffer and incubated for 10 min at 37°C with vehicle (retracted) or 10 μM calcium ionophore (protruded; A23187; Cayman Chemicals). 4 μl of the extracellular parasites were pipetted onto a glow-discharged (30 seconds at −30 mA) copper R2/2 holey carbon grid (Quantifoil Micro Tools GmbH, Jena, Germany). Samples were back-blotted with a Whatman filter paper (grade 1) for 3-4 s to remove excess liquid, then the grid was quickly plunge frozen into liquid ethane using a homemade plunge freezer. For detergent-extracted samples, membranes were extracted by addition of Triton-X-100 in HBSS for 3-4 min before washing briefly in HBSS and back-blotting, as above. Vitrified grids were mounted in notched Autogrids for cryo-FIB-milling (Thermo Fisher Scientific, MA, USA) and stored in liquid nitrogen until used.

### Cryo-FIB milling

Cryo-FIB milling was performed as previously described^67^. Briefly, Autogrids with vitrified *N. caninum* cells were loaded into a cryo-shuttle and transferred into an Aquilos dual-beam instrument (FIB/SEM; Thermo Fisher Scientific) equipped with a cryo-stage that is pre-cooled to minus 185 °C. Tile-set images of the grid were generated in SEM mode, and the cells suitable for cryo-FIB milling were targeted using the Maps software (Thermo Fisher Scientific). To protect the specimen and enhance conductivity, the sample surface was sputter-coated with platinum for 20 s at minus 30 mA current and then coated with a layer of organometallic platinum using the gas injection system pre-heated to 27°C for 5 s at a distance of 1 mm before milling^68^. Bulk milling was performed with a 30 kV gallium ion beam of 50 pA perpendicular to the grid on two side of a targeted cell. Then, the stage was tilted to 10°-18° between the EM grid and the gallium ion beam for lamella milling. For rough milling, the cell was milled with 30 kV gallium ion beams of 30 pA current, followed by 10 pA for polishing until the final lamella was 150 – 200 nm thick. The milling process was monitored by SEM imaging at 3 keV and 25 pA. A total of 178 lamella of *N. caninum* were milled over multiple sessions.

### Cryo-ET imaging

Cryo-FIB milled lamellae of vitrified apical regions of the parasites were imaged using a 300 keV Titan Krios transmission electron microscope (Thermo Fisher Scientific) equipped with a Bioquantum post-column energy filter (Gatan, Pleasanton, CA) used in zero-loss mode with a 20 eV slit width and a Volta Phase Plate with −0.5 μm defocus^69^. The microscope control software SerialEM was utilized to operate the Krios and collect tilt series from 56° to −56° in 2° increments using a dose-symmetric tilting scheme in the lose-dose mode^70,71^. Images were captured using a 5k × 6k K3 direct electron detection camera (Gatan) at a magnification of 26,000x (3.15 Å pixel size). Counting mode of the K3 camera was used, and for each tilt image 15 frames (0.04 s exposure time per frame, dose rate of ~28 e/pixel/s; frames were recorded in super-res mode and then binned by 2) were captured. The total electron dose per tilt series was limited to 100 e/Å^2^. In total, 167 tilt-series were collected from the cryo-FIB milled lamella of native parasites, but only about 10% of them contained the full or partial apical complex. 19 tilt-series were recorded of the apical region of detergent-treated (not cryo-FIB milled) parasites.

There are several factors that contribute to the low yield of the apical complex in our experiments. Firstly, we have plunge-frozen the intact parasites in a relatively thick (>1 μm) layer of ice. It is challenging to determine the apical vs. basal ends of the ice-embedded cells in the cryo-FIB milling instrument. Secondly, during the cryo-FIB milling step, the ice thickness was reduced from more than 1 μm to 150-200 nm, removing more than 80% of the volume. Thus, the probability of placing the lamella exactly in the region of the about 300 nm wide conoid is relatively low compared to accidentally milling the apical complex away during the thinning step. Finally, about 10~15% lamellae were damaged or surface-contaminated during the transfer step from the milling instrument to the TEM. Future application of fluorescence-guided cryo-FIB milling and autoloader systems for more direct transfer of FIB-milled samples into the TEM could address these issues and increase experiment throughput.

### Data processing and figure generation

The frames of each tilt series image were motion-corrected using MotionCor2 and then merged using the script extracted from the IMOD software package^72^ to generate the final tilt serial data set. Tilt series images were aligned either fiducial-less using patch tracking (800 × 800-pixel size) or using dark features as fiducials (*e.g*. granules from the sputter coat or embedded Gallium from the milling process) using the IMOD software package. Tomographic reconstructions were calculated using both weighted back-projection before subtomogram averaging, and simultaneous iterative reconstruction technique for visualizing raw tomogram data with higher contrast e.g. for particle picking. Of the recorded 167 tilt series of cryo-FIB milled native parasites, 125 tilt series were reconstructed for further inspection, and 20 of the reconstructed tomograms contained the apical complex (13 in the protruded, and 7 in the retracted state). Subtomograms that contain the conoid fiber, the ICMT or PCR repeats were extracted from the raw tomograms, aligned, and averaged with missing wedge compensation using the PEET program^73,74^. 1160 and 721 8-nm conoid fiber repeats were selected from the protruded and retracted native conoid tomograms, respectively, and 386 8-nm ICMT repeats were picked from the reconstructed tomograms of detergent-extracted samples. For the PCR average, 180 subtomograms from four tomograms (three protruded and one retracted conoid) were selected and initial motive lists with starting orientations were generated using the spikeInit functions in IMOD. Fourier shell correlations were calculated using the calcFSC function and plotted using the plotFSC function in IMOD. For the microneme tip, 25 subtomograms from three tomograms (two protruded and one retracted) were extracted, aligned and averaged. For visualization of raw tomographic slices, tomograms were denoised for improved clarity using either non-linear anisotropic diffusion or a weighted median filter (smooth filter) implemented in IMOD. Isosurface renderings and cellular segmentation with manual coloring were generated using the UCSF Chimera package software^75^, which was developed by the Resource for Biocomputing, Visualization, and Informatics at the University of California, San Francisco, with support from NIH P41-GM103311.

### Segmentation of the secretory organelles

Rhoptries, micronemes, and vesicles could be distinguished based on their distinctive morphologies. Rhoptries have a unique club shape and could be divided into two separate structural regions—an anterior tubular neck and a posterior bulb located deeper within the cell body. Micronemes are medium-sized, rod-like organelles and their interior displayed a darker electron density than other organelles. The vesicles that are closely associated with the ICMT or the apical plasma membrane are small (diameter < 50 nm), round or oval shaped, and their content appeared brighter (i.e. lower electron scattering properties) that the cytoplasm.

### Measurements

In order to measure the conoid dimensions, tomograms were rotated in the IMOD slicer windows, with the bottom plane oriented horizontally. The maximum distance in the longitudinal section across the center of the conoid was then used to determine the conoid’s dimensions as displayed in Supplemental Figure S5A. Three protruded and three retracted conoids were measured and compared. The lengths of the conoid fibers were measured as follows: After the conoid fiber was manually tracked using the IMOD program, the length of the conoid fibers and the spacing between adjacent conoid fibers were determined using the IMOD commands imodinfo and mtk, respectively. In total, 20 full-length conoid fibers from the protruded conoids and 12 from the retracted conoids were measured. To compare the APRs and PCRs in parasites with protruded or retracted conoid, we measured their diameters as follows: The tomograms were rotated in the IMOD slicer window until cross-sectional views of the PCRs and APRs could be saved (as seen in Supplemental Figure S5B,C); then the diameters of the rings were determined using the ImageJ circle tools. Measurements were made of three APRs and three PCRs from protruded conoids, and three APRs and two PCRs from retracted conoids. The distance between the conoid and the IMC were measured as follows: we positioned the center of the view in the IMOD slicer window at the apical end of the IMC and then rotated the tomogram around this center point in 3D to find the shortest distance between the conoid and the apical end of the IMC.

## Acknowledgments

We thank Daniel Stoddard for training and management of the cryo-electron microscopy facility at the University of Texas, Southwestern Medical Center, which is supported in part by the CPRIT Core Facility Support Award RP170644. This research was supported in part by the computational resources provided by the BioHPC supercomputing facility located in the Lyda Hill Department of Bioinformatics, UT Southwestern Medical Center.

## Authors contributions

M.L.R. and D.N. conceived the project; L.G. performed tomogram reconstruction, subtomogram averaging, data analysis, figure and movie preparation; W.J.O. performed cell culture, sample preparation and data analysis; K.C. performed cryo-preparation, cryo-ET data collection and initial image processing; E.R. performed cryo-FIB milling; M.L.R. performed data analysis and wrote the manuscript with help from L.G. and D.N.; D.N. performed data analysis and supervised the overall project.

## Data availability

The subtomogram average density maps of the *Neospora caninum* conoid fiber in the protruded, and retracted states, and a global average combining data from both states have been deposited in the Electron Microscopy Data Bank (EMDB) with accession code EMD-28247, EMD-28249, and EMD-26873, respectively. The subtomogram average density maps of the PCRs P1-P2, P3, and the composite map of P1-P2-P3 have been deposited in the Electron Microscopy Data Bank (EMDB) with accession code EMD-28231, EMD-28234, and EMD-28246, respectively. All other data needed to evaluate the conclusions in the paper are present in the paper and/or the Supplementary Materials. Additional data related to this paper may be requested from the corresponding authors.

## Figures

**Supplementary Materials for: Gui, et al.** Cryo-tomography reveals rigid-body motion and organization of apicomplexan invasion machinery

The PDF file includes:

Figs. S1 to S8

Other Supplementary Material for this manuscript includes the following:

**Movie S1**

## Supplemental Figures

**Figure S1.**
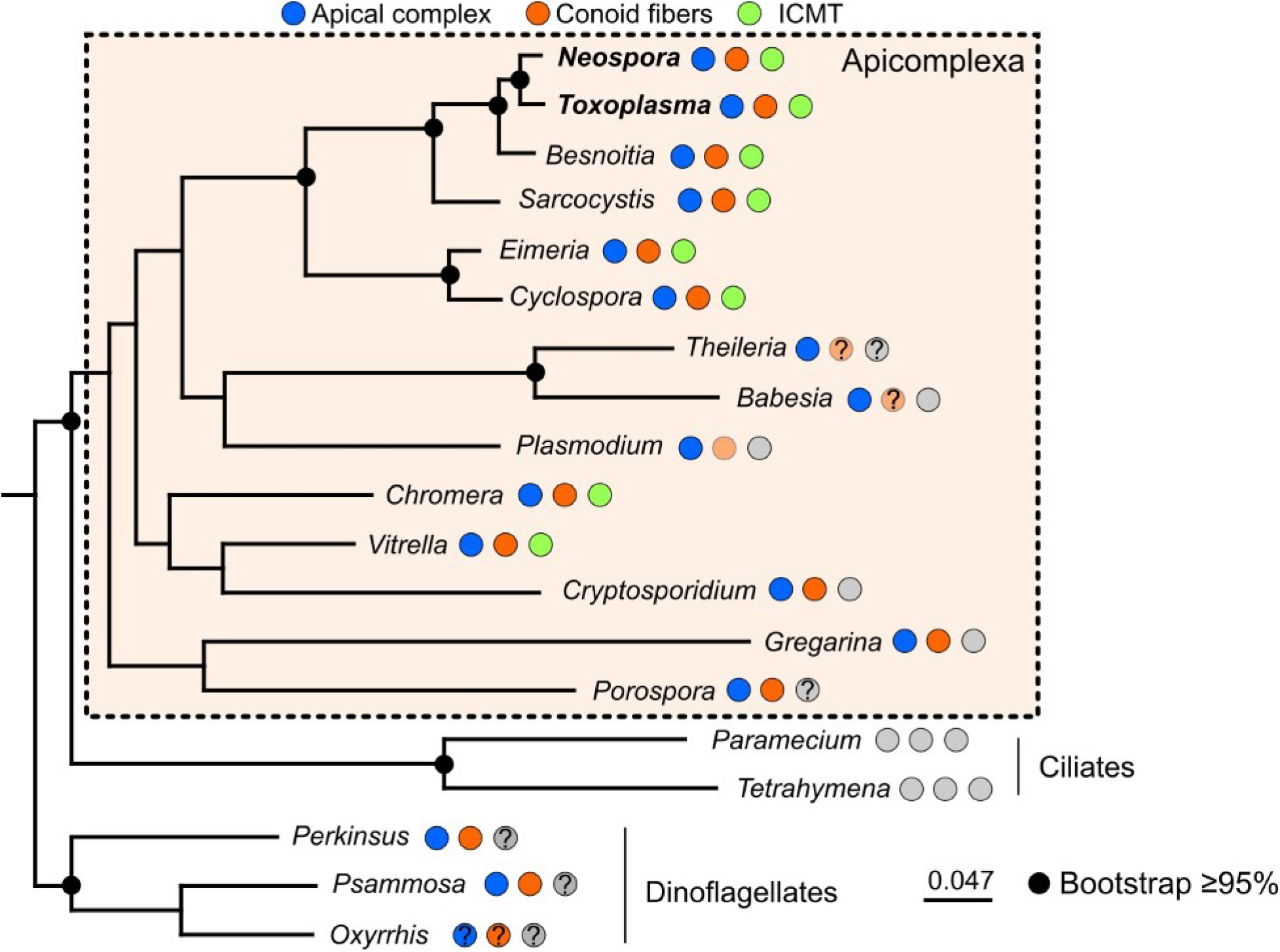
Phylogenetic comparison of Alveolate organisms. Phylogenetic tree of Apicomplexa and other alveolate organisms including representative ciliates and dinoflagellates. The presence of components of the apical complex is noted with colored circles next to each organism name. Note that the conoid tubulin fibers appear to be present in some, but not all, lifecycle stages of the Aconoidasida (*e.g. Plasmodium, Babesia*) that have a conoid structure (light orange circles).

**Figure S2.**
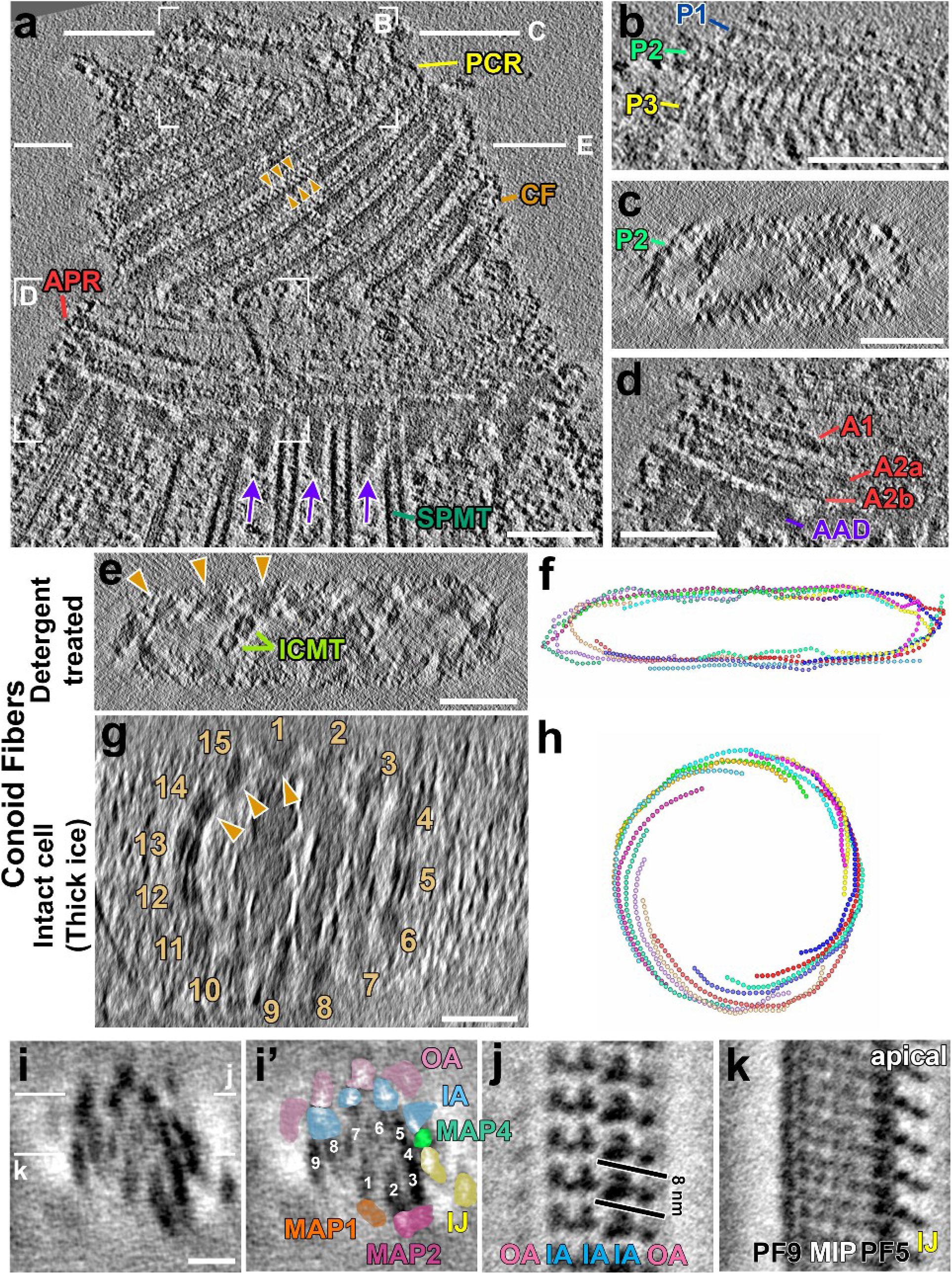
Detergent-extracted *Toxoplasma* exhibits a flattened structure in tomograms. **(A-D)** Tomographic slices of the apical complex of detergent-extracted tachyzoites in longitudinal (A, B, D) and cross-sectional orientation (C). White lines in (A) indicate the positions of the cross-sectional views shown in (C and E). Note the periodic connections between neighboring CFs (orange arrowheads in (A)). Other labels and coloring as for Figure 1. White boxes in (A) indicate the regions magnified and orientation-optimized in (B and D). (B) shows the detailed PCR structure with three layers (P1-P3) and repeating densities connecting the layers. A cross-sectional view of PCR-P2 (C) reveals severe flattening of the detergent-extracted sample. The APR-region is shown in (D) with at least three layers of APRs (A1, A2a, A2b) and a ring of amorphous APR-associated density (AAD). **(E-H)** Comparison of tomographic slices (E, G) and graphical models (F, H) of cross-sectional views of the apical complex in a detergent-extracted, severely flattened and distorted *Toxoplasma* cell (E, F), and in a native, un-flattened *N. caninum* cell that was embedded in a relatively thick ice layer and cryo-FIB milled to avoid cell flattening (G, H). Viewed from the apical end; dots/model points in (F, H) indicate repeat units of the CFs (orange arrowheads/1-15 in (E, G)). **(I-K)** Cross-sectional (I: original, I’: pseudo-colored) and longitudinal (J, K) tomographic slices through the subtomogram averaged 8-nm repeats of the conoid fibers in detergent-extracted apical complexes. Note that the resolution is relatively high in the longitudinal direction (J, K), but in cross-section (I), elongation due to the missing wedge artifact is apparent (flattening results in orientation bias of the CFs). Labels and coloring as in Figure 4. Scale bars: 100 nm (in A-E and G); 10 nm (in I, J and K).

**Figure S3.**
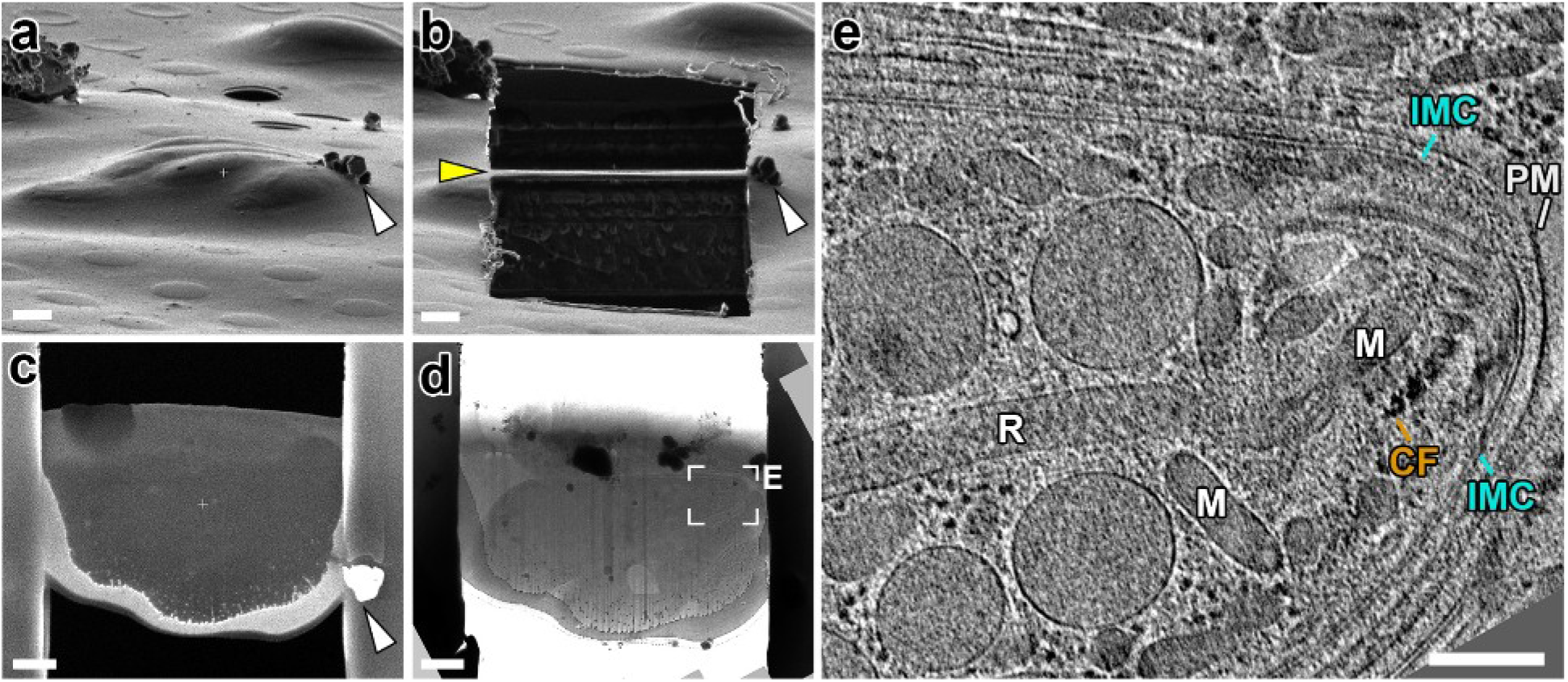
The cryo-FIB milling workflow to generate lamella of plunge-frozen *Neospora caninum* cells for cryo-ET imaging. **(A-C)** Ion beam images (A, B) and a scanning-EM image (C) show the same grid area with targeted *N. caninum* cells before (A) and after (B,C) cryo-FIB milling in side (A,B) and top (C) views. Yellow arrowhead in (B) indicates the about 200 nm thick cryo-FIB lamella. For reference, the same contamination particle (white arrowheads) is indicated in all ion beam and electron beam images. **(D)** A cryo-TEM image of the same lamella shown in (C) with two clearly visible *N. caninum* cells. A tilt series was recorded of the apical region (white box) of the top parasite. **(E)** Representative tomographic slice of the reconstructed apical complex (boxed in (D)), showing the conoid in the retracted state. Labels: CF, conoid fiber; IMC, inner membrane complex; M, microneme; PM, plasma membrane; R, rhoptry. Scale Bar, 1 μm (in A-D); 200 nm (in E).

**Figure S4.**
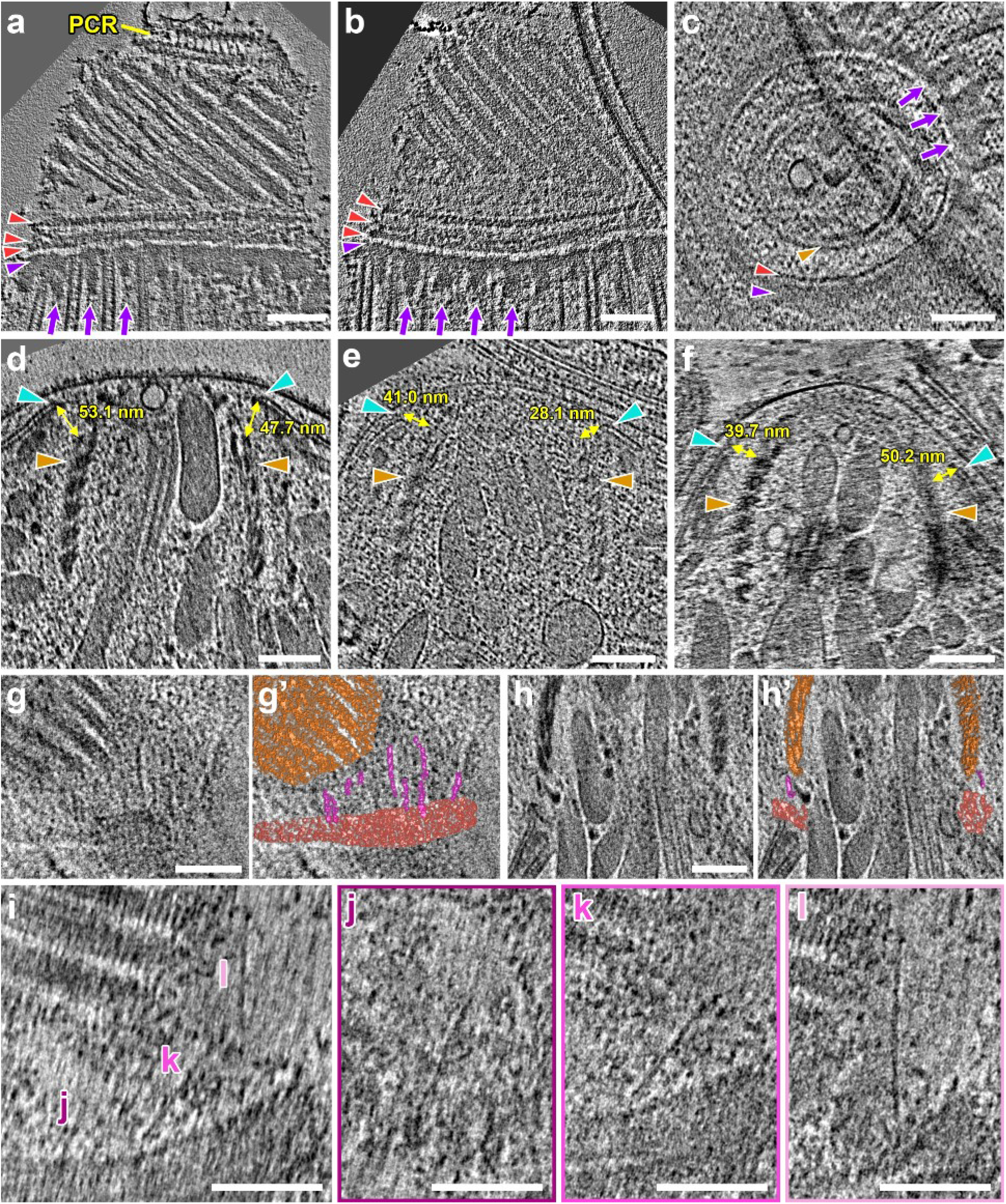
Flexible interactions between the APRs and the conoid. **(A, B)** Tomographic slices highlight the APRs of detergent-extracted *Toxoplasma* tachyzoites. **(C)** Cross-sectional tomographic slice of the APR in the partially retracted conoid from a cryo-FIB-milled *Neospora* tachyzoite. Red arrowhead indicates one of the annular rings of the APRs, purple arrowhead and arrows indicate the AAD ring and AAD projections, respectively. **(D-F)** Representative tomographic slices through cryo-FIB milled conoid complexes in the retracted state. Measurements mark the closest distance between the IMC apical edge (cyan arrowheads) and the conoid fibers (orange arrowheads). **(G-L)** Tomographic slices (G,H: original: G’,H’: pseudocolored) of protruded conoids show filamentous densities (magenta) with ~8 nm diameter and variable lengths that appear to connect the conoid fibers (orange) to the APRs (red). Note that the orientation of these filaments slightly varies within the 3D reconstructions, therefore it is impossible to capture all connections in full length in a single tomographic 2D slice (see Figure 2G for example 3D rendering). In order to clearly visualize three exemplary filaments that are partially visible in (I; marked with j-l), multiple slices at slightly different z-depths and 3D orientations are shown in (J-L). Scale bars: 100 nm.

**Figure S5.**
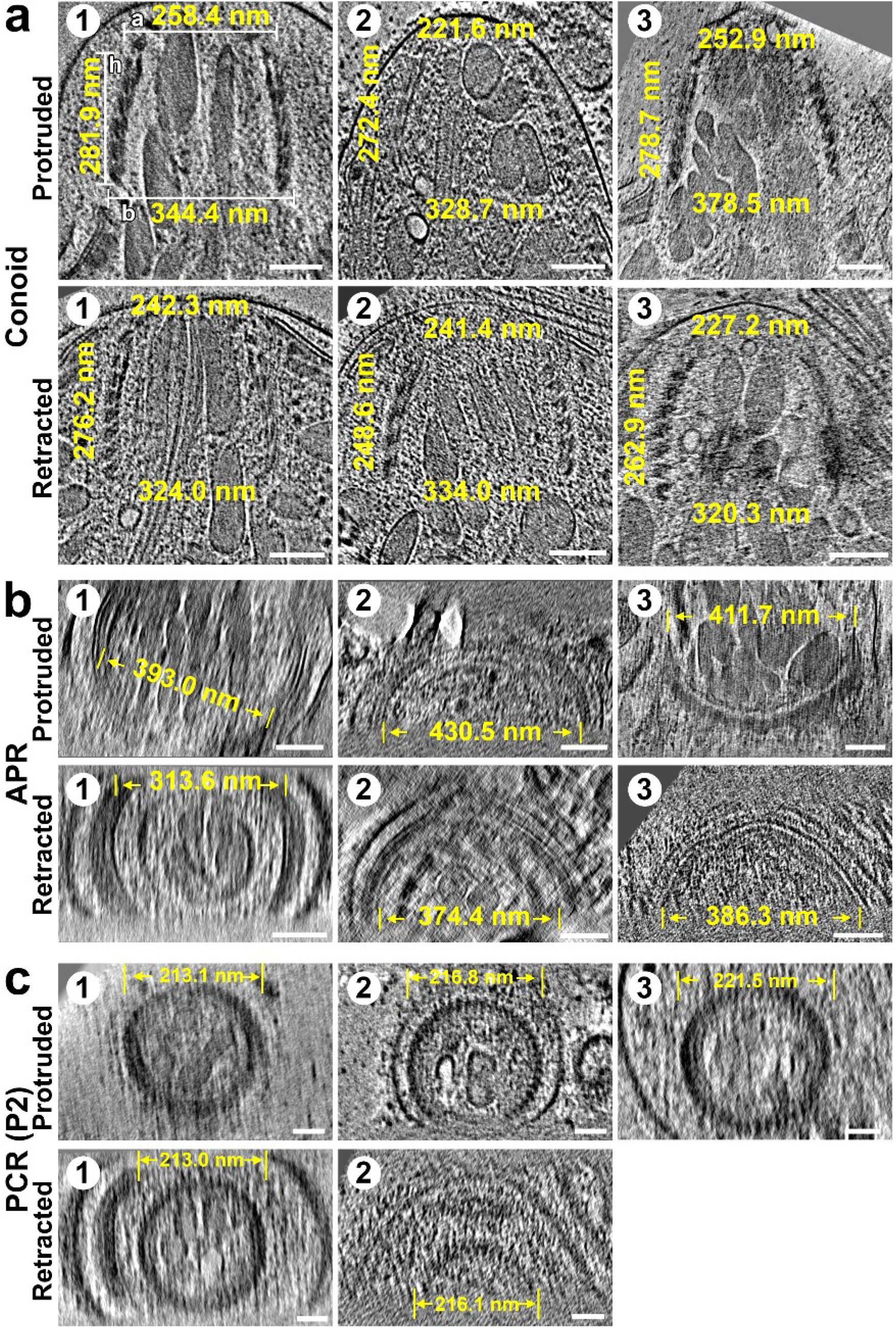
Image gallery of the conoid complex, APRs and PCRs in the protruded and retracted states of *N. caninum*. **(A)** Tomographic slices of the cryo-FIB-milled apical complex in the protruded and retracted states. Measurements indicate the apical diameter (a), the basal diameter (b) and the height (h) of the conoid structure (corresponds to analyses shown in Figure 3E). **(B, C)** Cross-sectional tomographic slices of apical complexes show the diameters of the APRs (B) and the P2-PCR (C) in the protruded and retracted states. Scale bars: 100 nm (A and B); 50 nm (C).

**Figure S6.**
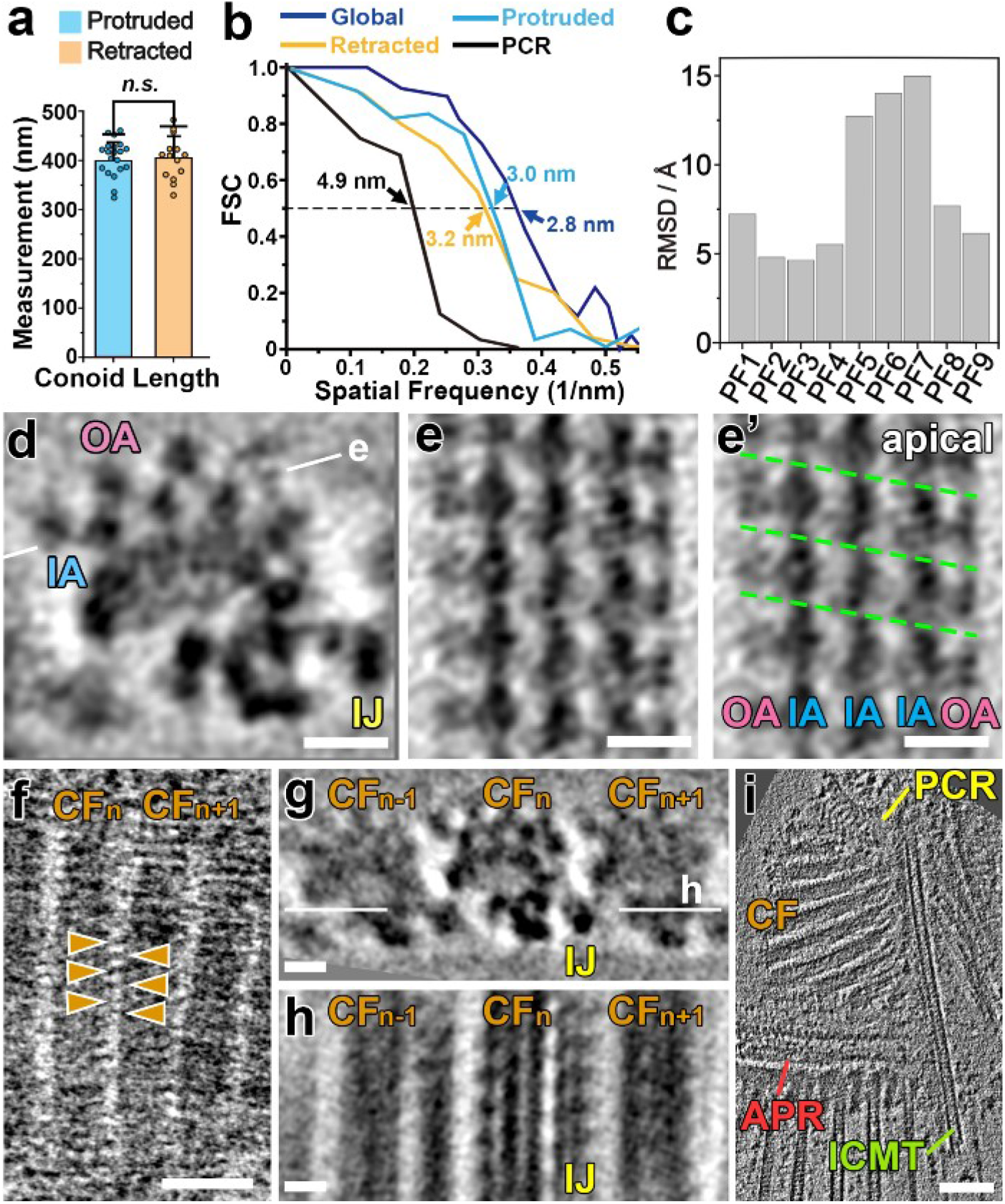
Structural features of the subtomogram averages of the conoid fibers from cryo-FIB milled *Neospora* cells (A-H), and the ICMT of a detergent-extracted *Toxoplasma* tachyzoite (I). **(A)** The lengths of the conoid fibers are unchanged between the protruded and retracted states. Statistical significance was calculated by Student’s t-test. n.s., not significant (p > 0.05). **(B)** Using the Fourier shell correlation method, resolution estimates of the global*, protruded, and retracted average of the conoid fibers were measured at the PF4 region, whereas the resolution of the PCR average was estimated at the P2 ring (* combining tomograms from both protruded and retracted states). Dashed line and values indicate where the FSC curve intersects with the 0.5 criteria. **(C)** RMSD values comparing the relative position and orientation of each protofilament in the pseudo-atomic models after fitting tubulin dimers to the averaged conoid fiber in the retracted and protruded states (see corresponding Figure 4E). **(D-E’)** Crosssectional (D) and longitudinal (E: original, E’: annotated) tomographic slices of the global subtomogram average of the conoid fibers show the pitch (indicated by green lines) along neighboring MAPs (E, E’). White lines in (D) indicate the positions of the longitudinal view in (E). **(F-H)** Tomographic slices through the conoid filaments in longitudinal orientation (F) show ~8-nm-spaced linkages (orange arrowheads) between adjacent conoid fibers (CF_n-1_, CF_n_ and CF_n+1_). In cross-sectional (G) and longitudinal (H) tomographic slices of the global subtomogram average of the conoid fibers (G, H; depicted are three neighboring conoid fibers) these linkages can be seen as the inner junction (IJ) MAPs between PF3 of conoid fibers CF_n_ and PF9 of the neighboring CF_n+1_. **(I)** Longitudinal tomographic slice of the apical complex of a detergent-extracted *Toxoplasma* tachyzoite shows the ICMT extending from the conoid apex passed the APRs. Scale bars: 10 nm (in D-E, G-H); 50 nm (in F); 100 nm (in I).

**Figure S7.**
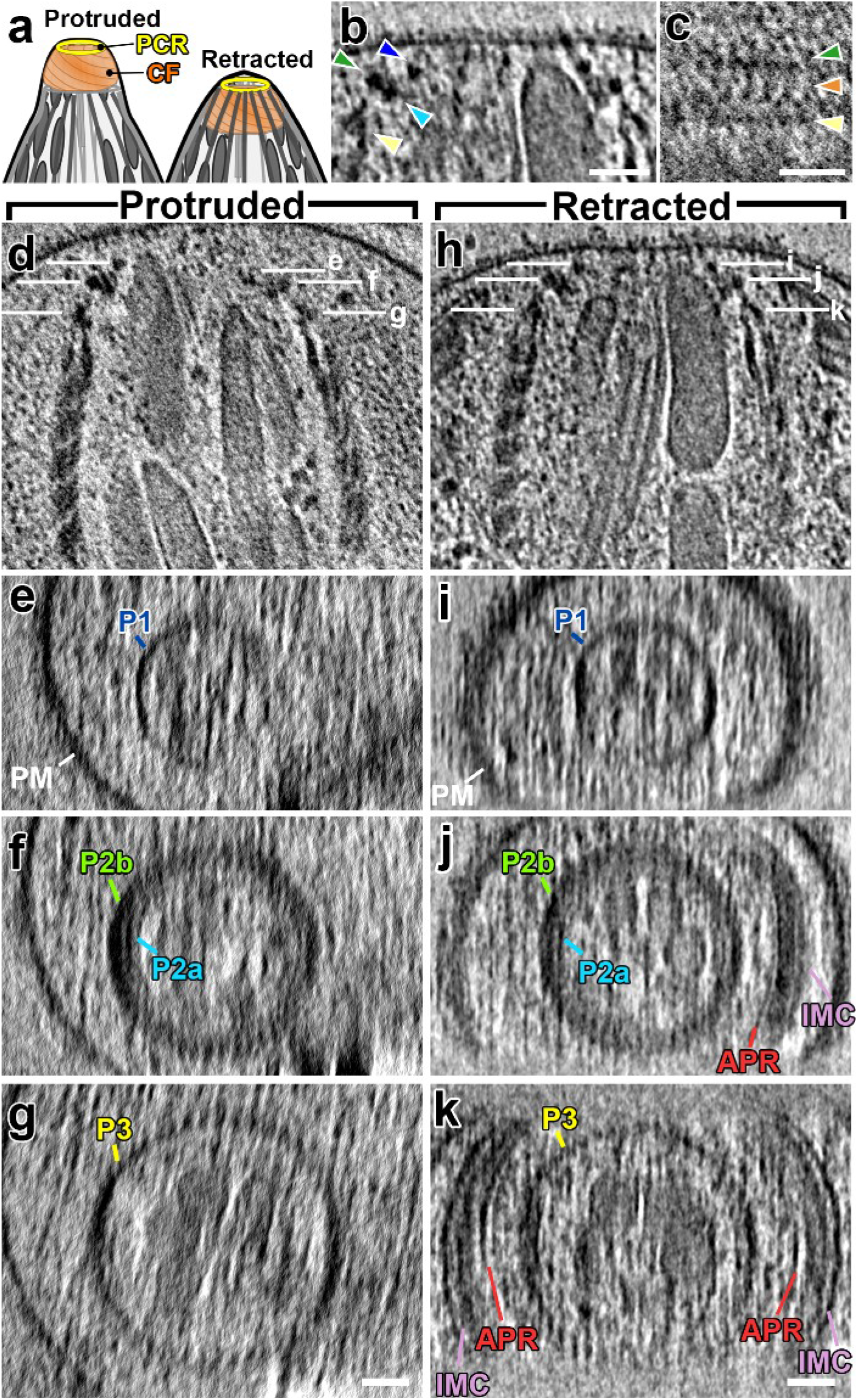
The PCR complex consists of three layers of rings. **(A)** Cartoon highlighting the relative positions of the PCRs and CFs in the apical complex. **(B, C)** Tomographic slices through the top of the apical complex in longitudinal (B) and tangential (C) orientations reveal different components of the PCRs, including the apical P1 (highlighted by blue arrowheads), P2a (cyan), P2b (green), P3 (yellow) and the linker (orange) between P2b and P3. **(D-K)** Longitudinal (D, H) and cross-sectional tomographic slices (E-G, I-K) through the top of the cryo-FIB-milled *N. caninum* apical complex in either protruded (D-G) or retracted states (H-K) reveal that the PCR complex is composed of at least three distinguishable annular structures that we name from apical to basal: P1 (blue), P2 – with an interior P2a (cyan) and an exterior P2b (green), and P3 (yellow). White lines in (D and H) indicate the positions of the corresponding cross-sectional views. Scale bars: 50 nm (in B, C, G, K, also valid for D-J).

**Figure S8.**
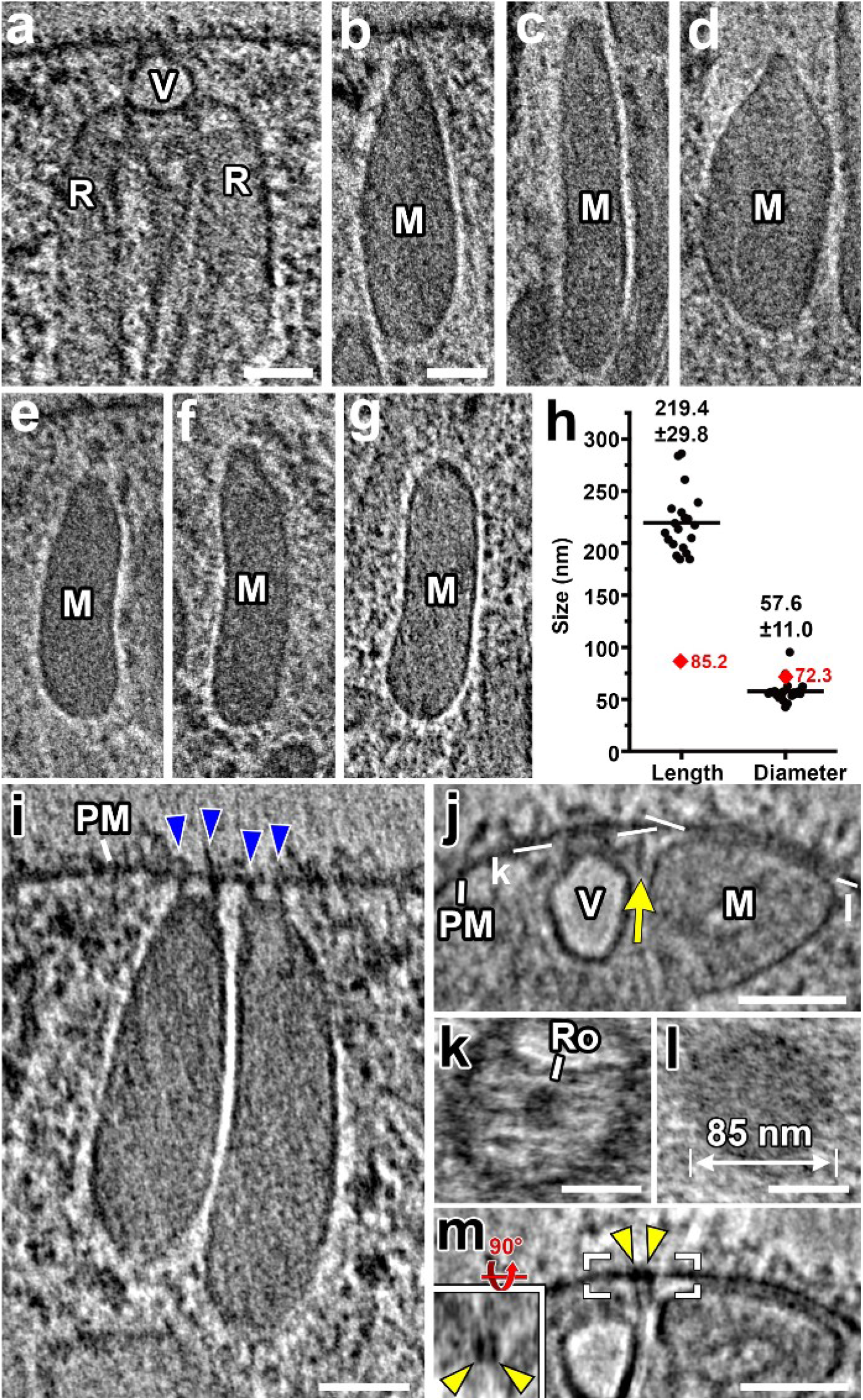
Structural details about the secretory organelles within the apical complex show that micronemes have a regular shape and size, and dock to the plasma membrane at sites distinct from but organized with the rhoptry-associated vesicle. **(A)** Tomographic slice displaying a membrane-docked apical vesicle (V) that associates with the apical tips of 2 rhoptries (R). **(B-G)** Representative tomographic slices of micronemes in cryo-FIB-milled *Neospora* tachyzoites highlight the fairly consistent shapes and polar morphology of these organelles; apical is oriented towards the top. **(H)** Distribution of lengths and diameters of the micronemes from the cryo-ET reconstructions. Numbers are averages ± standard deviations (n=20 micronemes). Red diamonds are measurements from the PM-docked microneme in Figure 7J, K, and Supplemental Figure S8J-M. Note that the length of this microneme is significantly less than the mean. **(I)** A tomographic slice showing two micronemes that connect to the plasma membrane (PM) through linkers (blue arrowheads), projecting from the corners of their flattened apical cap. **(J-M)** Tomographic slices provide side (J, M) and top views (K-L) of a microneme (M) and a rhoptry-associated vesicle (V) that are docked to the plasma membrane (PM) side-by-side (see also Figure 7J, K). The docking sites are distinct, i.e. the vesicle docks through the rosette (Ro in K) and the microneme via a contact site in a circular region of ~85 nm in diameter (L) to the plasma membrane. Despite the special separation, the location of docking appears to be organized in close proximity, likely through a long ridge (yellow arrow in J) that runs between the secretory organelles, and is tethered to a bi-partite anchor in the plasma membrane (yellow arrowheads in M and inset); the (M inset) shows the boxed area in (M) with the membrane anchor in top view. Scale bars: 50 nm (A, B, I-M; scale bar in B, also valid for C-G).

## References

1. Long, S., Anthony, B., Drewry, L. L. & Sibley, L. D. A conserved ankyrin repeat-containing protein regulates conoid stability, motility and cell invasion in Toxoplasma gondii. Nat. Commun. 8, 2236 (2017).

2. Long, S. et al. Calmodulin-like proteins localized to the conoid regulate motility and cell invasion by Toxoplasma gondii. PLoS Pathog. 13, e1006379 (2017).

3. Tosetti, N., Dos Santos Pacheco, N., Soldati-Favre, D. & Jacot, D. Three F-actin assembly centers regulate organelle inheritance, cell-cell communication and motility in Toxoplasma gondii. eLife 8, (2019).

4. O’Shaughnessy, W. J., Hu, X., Beraki, T., McDougal, M. & Reese, M. L. Loss of a conserved MAPK causes catastrophic failure in assembly of a specialized cilium-like structure in Toxoplasma gondii. Mol. Biol. Cell 31, 881–888 (2020).

5. Hu, K., Roos, D. S. & Murray, J. M. A novel polymer of tubulin forms the conoid of Toxoplasma gondii. J. Cell Biol. 156, 1039–1050 (2002).

6. Mondragon, R. & Frixione, E. Ca(2+)-dependence of conoid extrusion in Toxoplasma gondii tachyzoites. J. Eukaryot. Microbiol. 43, 120–127 (1996).

7. Carruthers, V. B. & Sibley, L. D. Mobilization of intracellular calcium stimulates microneme discharge in Toxoplasma gondii. Mol. Microbiol. 31, 421–428 (1999).

8. Del Carmen, M. G., Mondragón, M., González, S. & Mondragón, R. Induction and regulation of conoid extrusion in Toxoplasma gondii. Cell. Microbiol. 11, 967–982 (2009).

9. Hu, K. et al. Cytoskeletal components of an invasion machine--the apical complex of Toxoplasma gondii. PLoS Pathog. 2, e13 (2006).

10. Francia, M. E. et al. Cell division in Apicomplexan parasites is organized by a homolog of the striated rootlet fiber of algal flagella. PLoS Biol. 10, e1001444 (2012).

11. de Leon, J. C. et al. A SAS-6-like protein suggests that the Toxoplasma conoid complex evolved from flagellar components. Eukaryot. Cell 12, 1009–1019 (2013).

12. Wall, R. J. et al. SAS6-like protein in Plasmodium indicates that conoid-associated apical complex proteins persist in invasive stages within the mosquito vector. Sci. Rep. 6, 28604 (2016).

13. Bertiaux, E. et al. Expansion microscopy provides new insights into the cytoskeleton of malaria parasites including the conservation of a conoid. PLoS Biol. 19, e3001020 (2021).

14. Koreny, L. et al. Molecular characterization of the conoid complex in Toxoplasma reveals its conservation in all apicomplexans, including Plasmodium species. PLoS Biol. 19, e3001081 (2021).

15. Portman, N., Foster, C., Walker, G. & Šlapeta, J. Evidence of intraflagellar transport and apical complex formation in a free-living relative of the apicomplexa. Eukaryot. Cell 13, 10–20 (2014).

16. Okamoto, N. & Keeling, P. J. The 3D structure of the apical complex and association with the flagellar apparatus revealed by serial TEM tomography in Psammosa pacifica, a distant relative of the Apicomplexa. PloS One 9, e84653 (2014).

17. Azevedo, C. Fine structure of Perkinsus atlanticus n. sp. (Apicomplexa, Perkinsea) parasite of the clam Ruditapes decussatus from Portugal. J. Parasitol. 75, 627–635 (1989).

18. Satir, B. H. & Wissig, S. L. Alveolar sacs of Tetrahymena: ultrastructural characteristics and similarities to subsurface cisterns of muscle and nerve. J. Cell Sci. 55, 13–33 (1982).

19. Mann, T. & Beckers, C. Characterization of the subpellicular network, a filamentous membrane skeletal component in the parasite Toxoplasma gondii. Mol. Biochem. Parasitol. 115, 257–268 (2001).

20. Anderson-White, B. R. et al. A family of intermediate filament-like proteins is sequentially assembled into the cytoskeleton of Toxoplasma gondii. Cell. Microbiol. 13, 18–31 (2011).

21. Ouologuem, D. T. & Roos, D. S. Dynamics of the Toxoplasma gondii inner membrane complex. J. Cell Sci. 127, 3320–3330 (2014).

22. Ferreira, J. L. et al. The Dynamic Roles of the Inner Membrane Complex in the Multiple Stages of the Malaria Parasite. Front. Cell. Infect. Microbiol. 10, 611801 (2020).

23. Boothroyd, J. C. & Dubremetz, J.-F. Kiss and spit: the dual roles of Toxoplasma rhoptries. Nat. Rev. Microbiol. 6, 79–88 (2008).

24. Besteiro, S., Dubremetz, J. & Lebrun, M. The moving junction of apicomplexan parasites: a key structure for invasion. Cell Microbiol. 13, 797–805 (2011).

25. Santos, J. M. & Soldati-Favre, D. Invasion factors are coupled to key signalling events leading to the establishment of infection in apicomplexan parasites. Cell. Microbiol. 13, 787–796 (2011).

26. Aquilini, E. et al. An Alveolata secretory machinery adapted to parasite host cell invasion. Nat. Microbiol. 6, 425–434 (2021).

27. Plattner, H. et al. Genetic dissection of the final exocytosis steps in Paramecium tetraurelia cells: cytochemical determination of Ca2+-ATPase activity over performed exocytosis sites. J. Cell Sci. 46, 17–40 (1980).

28. McIntosh, R., Nicastro, D. & Mastronarde, D. New views of cells in 3D: an introduction to electron tomography. Trends Cell Biol. 15, 43–51 (2005).

29. Medalia, O. et al. Macromolecular architecture in eukaryotic cells visualized by cryoelectron tomography. Science 298, 1209–1213 (2002).

30. Lin, J. & Nicastro, D. Asymmetric distribution and spatial switching of dynein activity generates ciliary motility. Science 360, eaar1968 (2018).

31. Cyrklaff, M. et al. Cryoelectron tomography reveals periodic material at the inner side of subpellicular microtubules in apicomplexan parasites. J. Exp. Med. 204, 1281–1287 (2007).

32. Kudryashev, M. et al. Structural basis for chirality and directional motility of Plasmodium sporozoites. Cell. Microbiol. 14, 1757–1768 (2012).

33. Al-Amoudi, A., Studer, D. & Dubochet, J. Cutting artefacts and cutting process in vitreous sections for cryo-electron microscopy. J. Struct. Biol. 150, 109–121 (2005).

34. Marko, M., Hsieh, C., Schalek, R., Frank, J. & Mannella, C. Focused-ion-beam thinning of frozen- hydrated biological specimens for cryo-electron microscopy. Nat. Methods 4, 215–217 (2007).

35. Sun, S. Y. et al. Cryo-ET of Toxoplasma parasites gives subnanometer insight into tubulin-based structures. Proc. Natl. Acad. Sci. U. S. A. 119, e2111661119 (2022).

36. Nagayasu, E., Hwang, Y.-C., Liu, J., Murray, J. M. & Hu, K. Loss of a doublecortin (DCX)-domain protein causes structural defects in a tubulin-based organelle of Toxoplasma gondii and impairs host-cell invasion. Mol. Biol. Cell 28, 411–428 (2017).

37. Endo, T. & Yagita, K. Effect of extracellular ions on motility and cell entry in Toxoplasma gondii. J. Protozool. 37, 133–138 (1990).

38. Wang, X. et al. Cryo-EM structure of cortical microtubules from human parasite Toxoplasma gondii identifies their microtubule inner proteins. Nat. Commun. 12, 3065 (2021).

39. Leung, J. M. et al. Stability and function of a putative microtubule-organizing center in the human parasite Toxoplasma gondii. Mol. Biol. Cell 28, 1361–1378 (2017).

40. Tosetti, N. et al. Essential function of the alveolin network in the subpellicular microtubules and conoid assembly in Toxoplasma gondii. eLife 9, (2020).

41. Wetzel, D. M., Håkansson, S., Hu, K., Roos, D. & Sibley, L. D. Actin filament polymerization regulates gliding motility by apicomplexan parasites. Mol. Biol. Cell 14, 396–406 (2003).

42. Sibley, L. D., Hâkansson, S. & Carruthers, V. B. Gliding motility: an efficient mechanism for cell penetration. Curr. Biol. CB 8, R12–14 (1998).

43. Meissner, M., Schlüter, D. & Soldati, D. Role of Toxoplasma gondii myosin A in powering parasite gliding and host cell invasion. Science 298, 837–840 (2002).

44. Goddard, T. D., Huang, C. C. & Ferrin, T. E. Visualizing density maps with UCSF Chimera. J. Struct. Biol. 157, 281–287 (2007).

45. Mandelkow, E. M., Schultheiss, R., Rapp, R., Müller, M. & Mandelkow, E. On the surface lattice of microtubules: helix starts, protofilament number, seam, and handedness. J. Cell Biol. 102, 1067–1073 (1986).

46. Paredes-Santos, T. C., de Souza, W. & Attias, M. Dynamics and 3D organization of secretory organelles of Toxoplasma gondii. J. Struct. Biol. 177, 420–430 (2012).

47. Nichols, B. A. & Chiappino, M. L. Cytoskeleton of Toxoplasma gondii. J. Protozool. 34, 217–226 (1987).

48. Carruthers, V. B. & Sibley, L. D. Sequential protein secretion from three distinct organelles of Toxoplasma gondii accompanies invasion of human fibroblasts. Eur. J. Cell Biol. 73, 114–123 (1997).

49. Heaslip, A. T., Ems-McClung, S. C. & Hu, K. TgICMAP1 is a novel microtubule binding protein in Toxoplasma gondii. PloS One 4, e7406 (2009).

50. Coleman, B. I. et al. A Member of the Ferlin Calcium Sensor Family Is Essential for Toxoplasma gondii Rhoptry Secretion. mBio 9, (2018).

51. Mageswaran, S. K. et al. In situ ultrastructures of two evolutionarily distant apicomplexan rhoptry secretion systems. Nat. Commun. 12, 4983 (2021).

52. Bouchet-Marquis, C. et al. Visualization of cell microtubules in their native state. Biol. Cell 99, 45–53 (2007).

53. Morrissette, N. & Gubbels, M.-J. Chapter 13 - The Toxoplasma Cytoskeleton: Structures, Proteins and Processes. in Toxoplasma Gondii (Second Edition) (eds. Weiss, L. M. & Kim, K.) 455–503 (Academic Press, 2014). doi:10.1016/B978-0-12-396481-6.00013-1.

54. Lamarque, M. et al. The RON2-AMA1 interaction is a critical step in moving junction-dependent invasion by apicomplexan parasites. PLoS Pathog. (2011) doi:https://doi.org/10.1371/journal.ppat.1001276.

55. Tyler, J. S. & Boothroyd, J. C. The C-terminus of Toxoplasma RON2 provides the crucial link between AMA1 and the host-associated invasion complex. PLoS Pathog. 7, e1001282 (2011).

56. Aikawa, M. Ultrastructure of the pellicular complex of Plasmodium fallax. J. Cell Biol. 35, 103–113 (1967).

57. de Souza, W. Fine structure of the conoid of Toxoplasma gondii. Rev. Inst. Med. Trop. Sao Paulo 16, 32–38 (1974).

58. Monteiro, V. G., de Melo, E. J., Attias, M. & de Souza, W. Morphological changes during conoid extrusion in Toxoplasma gondii tachyzoites treated with calcium ionophore. J. Struct. Biol. 136, 181–189 (2001).

59. Leung, J. M. et al. A doublecortin-domain protein of Toxoplasma and its orthologues bind to and modify the structure and organization of tubulin polymers. BMC Mol. Cell Biol. 21, (2020).

60. Katris, N. J. et al. The apical complex provides a regulated gateway for secretion of invasion factors in Toxoplasma. PLoS Pathog. 10, e1004074 (2014).

61. Dos Santos Pacheco, N. et al. Conoid extrusion regulates glideosome assembly to control motility and invasion in Apicomplexa. Nat. Microbiol. (2022) doi:10.1038/s41564-022-01212-x.

62. Jacot, D., Daher, W. & Soldati-Favre, D. Toxoplasma gondii myosin F, an essential motor for centrosomes positioning and apicoplast inheritance. EMBO J. 32, 1702–1716 (2013).

63. Periz, J. et al. A highly dynamic F-actin network regulates transport and recycling of micronemes in Toxoplasma gondii vacuoles. Nat. Commun. 10, 4183 (2019).

64. Carmeille, R., Schiano Lomoriello, P., Devarakonda, P. M., Kellermeier, J. A. & Heaslip, A. T. Actin and an unconventional myosin motor, TgMyoF, control the organization and dynamics of the endomembrane network in Toxoplasma gondii. PLoS Pathog. 17, e1008787 (2021).

65. Håkansson, S., Charron, A. J. & Sibley, L. D. Toxoplasma evacuoles: a two-step process of secretion and fusion forms the parasitophorous vacuole. EMBO J. 20, 3132–3144 (2001).

66. Stamatakis, A. RAxML version 8: a tool for phylogenetic analysis and post-analysis of large phylogenies. Bioinforma. Oxf. Engl. 30, 1312–1313 (2014).

67. Hariri, H. et al. Mdm1 maintains endoplasmic reticulum homeostasis by spatially regulating lipid droplet biogenesis. J. Cell Biol. 218, 1319–1334 (2019).

68. Schaffer, M. et al. Optimized cryo-focused ion beam sample preparation aimed at in situ structural studies of membrane proteins. J. Struct. Biol. 197, 73–82 (2017).

69. Danev, R., Tegunov, D. & Baumeister, W. Using the Volta phase plate with defocus for cryo-EM single particle analysis. eLife 6, e23006 (2017).

70. Mastronarde, D. N. Automated electron microscope tomography using robust prediction of specimen movements. J. Struct. Biol. 152, 36–51 (2005).

71. Hagen, W. J. H., Wan, W. & Briggs, J. A. G. Implementation of a cryo-electron tomography tiltscheme optimized for high resolution subtomogram averaging. J. Struct. Biol. 197, 191–198 (2017).

72. Kremer, J. R., Mastronarde, D. N. & McIntosh, J. R. Computer visualization of three-dimensional image data using IMOD. J. Struct. Biol. 116, 71–76 (1996).

73. Nicastro, D. et al. The molecular architecture of axonemes revealed by cryoelectron tomography. Science 313, 944–948 (2006).

74. Heumann, J. M., Hoenger, A. & Mastronarde, D. N. Clustering and variance maps for cryo-electron tomography using wedge-masked differences. J. Struct. Biol. 175, 288–299 (2011).

75. Pettersen, E. F. et al. UCSF Chimera--a visualization system for exploratory research and analysis. J. Comput. Chem. 25, 1605–1612 (2004).

